# Psilocybin-enhanced fear extinction linked to bidirectional modulation of cortical ensembles

**DOI:** 10.1101/2024.02.04.578811

**Authors:** Sophie A. Rogers, Elizabeth A. Heller, Gregory Corder

**Author notes:** Correspondence (G.C.).

## Abstract

The serotonin 2 receptor (5HT2R) agonist psilocybin displays rapid and persistent therapeutic efficacy across neuropsychiatric disorders characterized by cognitive inflexibility. However, the impact of psilocybin on patterns of neural activity underlying sustained changes in behavioral flexibility has not been characterized. To test the hypothesis that psilocybin enhances behavioral flexibility by altering activity in cortical neural ensembles, we performed longitudinal single-cell calcium imaging in the retrosplenial cortex across a five-day trace fear learning and extinction assay. A single dose of psilocybin induced ensemble turnover between fear learning and extinction days while oppositely modulating activity in fearand extinctionactive neurons. The acute suppression of fear-active neurons and delayed recruitment of extinction-active neurons were predictive of psilocybin-enhanced fear extinction. A computational model revealed that acute inhibition of fear-active neurons by psilocybin is sufficient to explain its neural and behavioral effects days later. These results align with our hypothesis and introduce a new mechanism involving the suppression of fear-active populations in the retrosplenial cortex.

## Introduction

Neuropsychiatric disorders characterized by inflexible associative learning, such as depression, anxiety, substance use-disorders, and post-traumatic stress disorder, affect over 350 million people worldwide^1^. Serotonergic psychedelics, including psilocybin, demonstrate remarkable transdiagnostic potential across these disorders^2^. After only a single dose, many patients report long-lasting improvements in depression and SUDs, as well as overall well-being *for up to a year*—a time-span implicating the involvement of cortically mediated long-term memory^3–6^. Therapeutic-like effects also have been observed in rodent models in many behavioral studies^7–15^, enabling the study of the neural mechanisms of psilocybin-enhanced mental health outcomes in mice.

Psilocybin is a naturally occurring compound found in hundreds of species of mushroom. Upon first pass metabolism, psilocybin is dephosphorylated into its active metabolite psilocin – a potent serotonin receptor agonist^16,17^. While psilocybin’s subjective effects tend to be accompanied by feelings of extreme “bliss”, “unity”, and “meaningfulness”^2,18^, in a subset of patients, psilocybin can induce anxiogenic and even traumatic experiences, in some cases associated with long-term psychosis and suicidal ideation^19-24^. To develop safe therapies with minimal adverse side-effects, it is critical to identify the relevant neural subpopulations differentially modulated by psilocybin to produce long-lasting therapeutic effects.

Clinical researchers found that therapeutic effects of serotonergic psychedelics in humans are mediated by increased cognitive flexibility following drug experience, a finding recapitulated in rodent models^25-28^. Evidence from human, rodent, and molecular research converges on the hypothesis that psilocybin generates highly plastic brain states conducive to modifying circuits that underlie inflexible, maladaptive behaviors via 5HT2R and TrkB activation^2,17,29-36^. Acute activation of cortical neurons by psychedelics induces synaptic AMPA receptor insertion, BDNF signaling, and consequent dendritic growth^32,35,37,38^. It is unknown how these molecular actions of psilocybin impact information processing in neural ensembles associated with aversive memories and maladaptive behavioral patterns.

The retrosplenial cortex (RSC) is one region where psilocybin may alter information processing in a manner sustaining enhanced behavioral flexibility. The RSC implements a variety of abstract functions^39^, including encoding and retrieval of episodic memory^40-51^; imagination of the future^39^; value and context encoding^44-48^; egocentric navigation and reasoning^49,52-54^; and ego dissolution under psychedelics^55^. Chemogenetically inhibiting RSC during reversal learning impairs performance after a rule switch, suggesting RSC activity is crucial for behavioral flexibility^48^. Replay of neural activation in the RSC is also involved in the production, consolidation, and generalization of fear engrams^51^. In another study, Wang et al., 2019 identified a previously silent ensemble recruited in RSC during contextual fear extinction, another form of behavioral flexibility^56^. When the authors optogenetically reactivated this ensemble after re-conditioning, extinction was reinstated, suggesting that excitatory plasticity in the RSC drives fear extinction. The increase in c-FOS-expressing neurons after extinction observed by Wang et al. was recently replicated and shown to be sex-independent^56,57^.

Several lines of evidence suggests that psilocybin’s effects could in part be mediated by changes in RSC activity. Psilocybin increases c-FOS expression throughout the cortex but idiosyncratically alters neural oscillations specifically in the RSC^58,59^. While 5HT2ARs are distributed throughout cortical L5 pyramidal neurons, the RSC is the only cortical region that also contains 5HT2CRs on pyramidal neurons^60,61^. In humans with depression, functional connectivity between the dorsal raphe nuclei and posterior cingulate regions homologous to rodent RSC is impaired^62^. Subsequent improvements in functional connectivity between the posterior cingulate and prefrontal cortex predict psilocybin-induced enhancements in cognitive flexibility^20^. Importantly, the RSC is involved in the retrieval of remote fear memories, positioning it as a potential substrate for psilocybin’s longer-lasting effects^62-66^.

To investigate the role of the RSC in the post-acute effects of psilocybin on behavioral flexibility, extinction of trace fear conditioning (TFC) was employed as the appropriate primary behavioral paradigm. TFC is a model of complex fear learning in rodents, in which a conditioned stimulus (CS) is followed by a trace period of 20-seconds preceding the shock. The trace period in TFC renders conditioning and extinction cortex-dependent, requiring protracted attention to associate temporally distant stimuli^68-70^. When the shock is omitted during extinction, animals must learn that it is now safe to reduce their freezing response or extinguish. In mice, Kwapis et al. found that TFC extinction depends on excitatory activity in the RSC^68^. Others have shown that optical, electrolytic, and pharmacological interventions in the RSC impact various kinds and stages of FC^45,49,50,55,63-71^. In a one-day paradigm, psilocybin administered 24 hours prior facilitated TFC extinction at low doses^72^. However, this study did not investigate the effect of psilocybin on long-term, consolidated fear memory, which is of translational interest.

To evaluate the hypothesis that psilocybin promotes behavioral flexibility by rapidly and persistently altering RSC ensembles associated with aversive memories, we investigated the effects of a single dose of psilocybin in a multi-day TFC extinction paradigm. We repeated this experiment in GCaMP8m-expressing, miniature microendoscope-implanted mice to measure singlecell calcium activity throughout the task. Using tensor component analysis (TCA)^73^, we identified ensembles driving RSC activity during different cognitive phases of the task acquisition, early extinction/fear recall, and late extinction. We confirmed our hypothesis that psilocybin accelerates and enhances the recruitment of an extinction-associated ensemble, particularly in drug-responsive animals. To our surprise, we found that fear extinction was also associated with an acute, robust suppression of fear-associated neurons during psilocybin administration. These effects on neural activity predicted extinction, and a computational model revealed that this inhibition is in fact sufficient to explain enhanced recruitment in extinction-associated neurons as well as behavioral variability. Taken together, these results support two mechanisms of psilocybin-enhanced fear extinction in the RSC, based on opposing forms of plasticity, which act in concert to reduce behavioral inflexibility.

## Results

### Psilocybin enhances Trace Fear Extinction (TFC) extinction in a responsive subpopulation of mice

Mice underwent a five-day TFC paradigm, with one Habituation, one Acquisition, and three Extinction Sessions (**Fig. 1a**). Freezing was measured in ezTrack^74^ as percent of time immobile during the trace period. Mice were administered psilocybin (1.0 mg/kg, i.p.) or saline 30 min before *Extinction 1*. This time-point was chosen as psilocybin-induced head twitches, the behavior taken as a proxy for the subjective effects in animals, peak around 15 min and last for up to 150 min^42^. Both groups froze significantly more during the first trace period of Extinction 1 than baseline, indicating successful TFC acquisition (**Supplementary Fig. 1e, Supplementary Table 1**). Psilocybin did not affect recall during Extinction 1, defined as freezing in the first half of the session (**Fig 1b, Supplementary Table 1**). Nonetheless, psilocybin mice reduced freezing more quickly during Extinction 1, indicating that psilocybin acutely accelerated fear extinction. (**Supplementary Fig 1a, Supplementary Table 1**) Psilocybin significantly reduced freezing during early Extinction 2, indicating enhanced extinction recall. By Extinction 3 there was no significant difference between groups, nor at a one-month follow-up (**Fig. 1b, Supplementary Fig. 1a,f, Supplementary Table 1**). Overall, there was no difference between males and females in either condition (**Supplementary Fig. 1b,c, Supplementary Table 1**).

**Figure 1.**
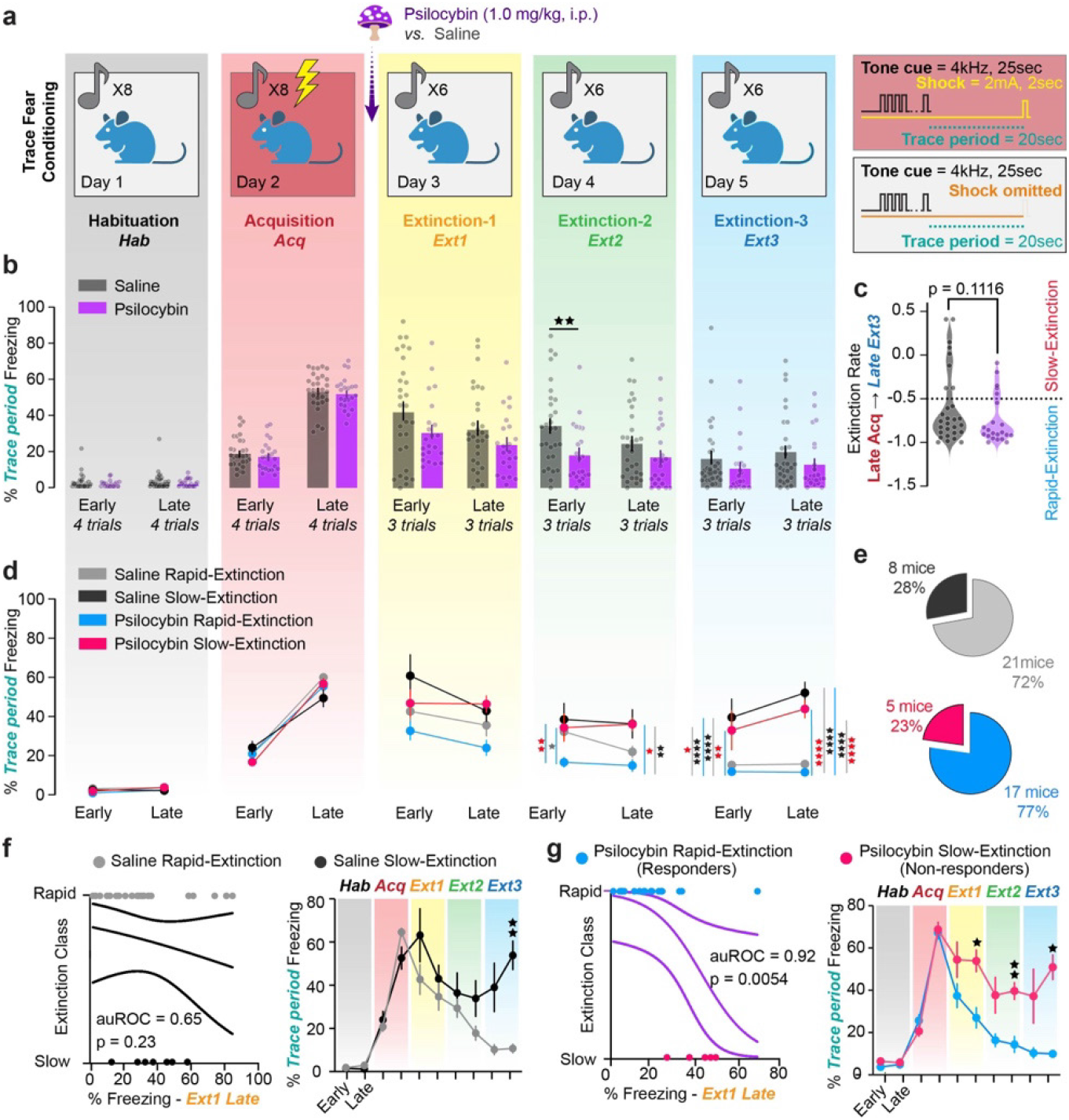
Psilocybin enhances TFC extinction in a responsive subpopulation of mice. **a**. Diagram of five-day TFC experiment. Right-hand panels depict conditioned and unconditioned one parameters. **b**. Average % time freezing during trace period in the first and last 3 trials of each day (“Early,” “Late” respectively) in saline and psilocybin-administered mice (black and purple respectively, n=25 each). Dots are individual animals. Two-Way ANOVA with Sidak multiple comparisons correction (Supp. Table 1, rows 1-5) **c**. Extinction rate calculated as the difference between freezing during late Acquisition and late Extinction 3 divided by freezing during late Acquisition. Red line indicates -50% threshold distinguishing rapidlyfrom slowly-extinguishing mice. Unpaired t-test. (Supp. Table 1, rows 6) **d**. Same as B; treatment groups subdivided into rapid-and slow-extinguishing mice (light colors, rapid; dark colors, slow). Two-Way ANOVA with Sidak multiple comparisons correction. (Supp. Table 1, rows 7-11) **e**. Pie charts describing breakdown of rapid-and slow-extinguishing mice within treatment groups. **f**. Left: Logistic regression predicting extinction rate based on % time freezing during early *Extinction 1* during acute drug treatment in saline-administered mice. Right: Direct comparison of % freezing over time between saline rapid-and slow-extinguishing mice. 2-Way ANOVA. (Supp. Table 1, rows 12-13) **g**. Same as F for psilocybin-administered mice. (Supp. Table 1, rows 14-15) Data are mean ± SEM. * p ≤ 0.05, ** p < 0.01, *** p < 0.001, **** p < 0.0001.

The extinction rate was calculated as the percent difference between freezing in late Acquisition and late Extinction 3. We chose this definition to circumvent the confounding variable of drug-induced changes in mobility during Extinction 1. Notably, there was a trend towards psilocybin-enhanced extinction rate, with a skewed distribution of extinction rates in psilocybin and saline groups (**Fig. 1c, Supplementary Table 1**). As extinction rate was one of our primary outcomes, mice that had extinguished >50% of their late Acquisition freezing by late Extinction 3 were classified as rapidly extinguishing and all others as slowly extinguishing.

Intriguingly, psilocybin-administered rapidly extinguishing mice reduced freezing more quickly than all other groups (**Fig. 1d, Supplementary Table 1**). This effect was greatest during early Extinction 2, suggesting psilocybin particularly enhanced recall of extinction memory. In contrast, freezing in slowly extinguishing mice was unaffected by treatment (**Fig. 1d, Supplementary Table 1**). There was the same proportion of rapid and slow extinguishing mice in each group (**Fig. 1e**).

To determine whether there was subpopulation of psilocybin non-responsive mice or whether it is always the case that mice that freeze more during recall extinguish slowly, we asked if freezing during the psilocybin’s acute effects in Extinction 1 would predict extinction class. (**Fig. 1f,g left, Supplementary Table 1**) Interestingly, the percent freezing during late Extinction 1 predicted classification as either rapidly or slowly extinguishing only if mice were administered psilocybin (auROC = 0.9176, p = 0.0054), but not saline (auROC = 0.6488, p = 0.2225). Indeed, freezing in psilocybin rapid-extinguishers was significantly lower than slow-extinguishers during acute psilocybin administration and consistently throughout the subsequent days. On the other hand, saline rapid-and slow-extinguishers are only differentiated at the timepoints used to define the classes, suggesting there were no pre-existing distinctions between the groups. Thus, we identified two classes of psilocybin-responsive versus non-responsive mice, hereon referred to as “responders” and “non-responders” respectively.

### Miniscope-implanted mice acquire and extinguish TFC

To explore the neurophysiological correlates of psilocybin-enhanced TFC extinction, single cell calcium activity was recorded in the RSC of saline and psilocybin mice. Mice were injected with AAV9-*hSyn*-GCaMP8m in the RSC and two weeks later implanted with a 1.0 mm diameter, gradient refractive index (GRIN) lens over the injection site (**Fig. 2a B**). After 2-5 weeks, mice were trained in the same TFC task (**Fig. 2c**). Calcium traces were extracted using CNMF-E in the Inscopix Data Processing Software API and post-processed (**Fig. 2d**). Across the entire TFC imaging protocol, 11-160 RSC neurons per animal (median = 46) were longitudinally registered across all days (Psilocybin responders = 460 total neurons; Psilocybin non-responders = 350 total neurons; Saline rapidly-extinguishing mice = 241 total neurons; Saline slowly-extinguishing mice = 116 neurons total; **Fig. 2e,g,h**). Miniscope placements were validated in all mice (**Supplementary Fig. 2**).

**Figure 2.**
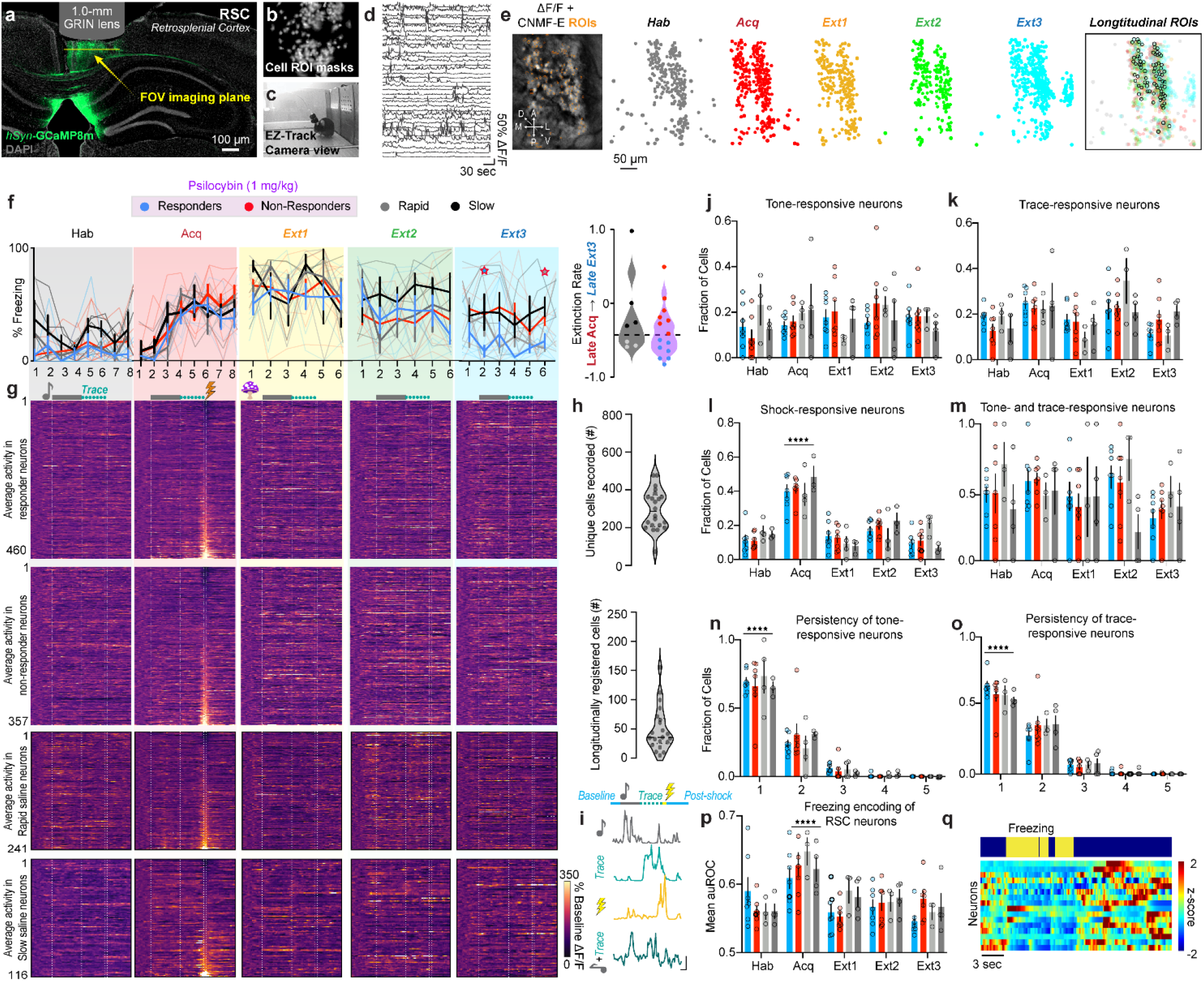
RSC neurons are modulated over TFC extinction in psilocybin- and saline-administered mice. **a**. Representative image of *AAV8-syn-GCaMP8m-WPRE* expression (green) and nuclei (grey) in RSC under GRIN lens tract. **b**. Cell masks of imaged neurons during one session from same mouse. **c**. Image of behavioral set-up during an extinction session. Example frame of freezing mouse. **d**. Example traces of neurons recorded during behavior in dF/F in same mouse. **e**. Left: Representative image from Ca^2+^-imaging video in the same mouse. Right: Cell masks of each recorded neuron in each session, overlayed with masks of longitudinally registered cells. **f**. Left: Percent of time freezing during each trial in responders (n=7 mice), non-responders (n=7 mice), and saline mice (n=7 mice). Two-Way ANOVA. (Supp. Table 1, rows 16-20) Right: Extinction rate. Unpaired t-test. (Supp. Table 1, row 21). **g**. Average activity in each neuron over all trials from each session, normalized to baseline before one onset and aligned to shock. Top: Responders (n=460 neurons), Top middle: Non-responders (n=357 neurons), Bottom middle: rapid saline (n=241 neurons), Bottom: slow saline (n=116 neurons). **h**. Top: Number of unique cells accepted over all sessions in each animals. Bottom: Number of longitudinally registered neurons in each animal. **i**. Example traces of tone-, trace-, shock, and tone+trace-responsive neurons (top to bottom). Vertical scale bar = 2dF/F, horizontal scale bar = 5 sec. **j**. Fraction of tone-responsive cells in each group over each day. Two-Way RM ANOVA. (Supp. Table 1, row 22) **k**. Fraction of trace-responsive cells in each group over each day. Two-Way RM ANOVA. (Supp. Table 1, row 23) **L**. Fraction of shock/omission-responsive cells in each group over each day. Two-Way RM ANOVA. (Supp. Table 1, row 24) **m**. Fraction of one-responsive neurons that are also trace-responsive cells in each group over each day. Two-Way RM ANOVA. (Supp. Table 1, row 25) **n**. Fraction of one-responsive cells that are one-responsive for 1-5 days in each animal. Two-Way RM ANOVA. (Supp. Table 1, row 26) **o**. Fraction of trace-responsive cells that are trace-responsive for 1-5 days in each animal. Two-Way RM ANOVA. (Supp. Table 1, row 27) **p**. Average freezing encoding of neurons in each group over each day (auROC, Two-Way ANOVA). (Supp. Table 1, row 28) **q**. Representative traces of freezing-encoding neurons in 1 animal sorted from greatest to least (bottom to top) auROC. Data are represented as mean ± SEM. * p ≤ 0.05, ** p < 0.01, *** p < 0.001, **** p < 0.0001.

All miniscope mice successfully acquired TFC and were subsequently split into psilocybin (1.0 mg/kg; i.p.; n=14 mice) and saline (n=7 mice; **Fig. 2f left, Supplementary Table 1**). Miniscope implanted mice extinguished less robustly than surgically naïve mice, indicating an impact either of head trauma or chronic stress post-implantation on TFC extinction (**Fig. 2f right, Supplementary Table 1**). Nonetheless, by Extinction 3, a subset of psilocybin responders emerged. Seven of fourteen psilocybin-treated mice and three of seven saline-treated mice extinguished their freezing by over 50% (**Fig. 2f, Supplementary Table 1**).

### RSC neurons are modulated over TFC training

To determine changes in the task-relevant response properties of RSC neurons, fractions of tone-, trace, shock-, and tone+trace-responsive neurons were measured each day (**Fig. 2i**). Fractions of toneor trace period-upregulated neurons were not significantly affected over time in any treatment group and in general varied between 10-30% of neurons (**Fig. 2j,k, Supplementary Table 1**). ∼40% of recorded RSC neurons were shock-responsive neurons (**Fig. 2l, Supplementary Table 1**). On average, ∼50% of tone-responsive neurons on a given day were also trace-responsive on the same day, suggesting a high degree of overlap of activated neurons between different periods in a trial (**Fig. 2m, Supplementary Table 1**). There was a large rate of turnover in tone- and trace-responsive neurons between days, with ∼75% of tone- and ∼60% trace-responsive neurons maintaining their responsiveness for only 1 day and about ∼25% and ∼30% respectively for 2 days across groups (**Fig. 2n,o, Supplementary Table 1**).

Similar proportions of all neurons recorded were responsive to various stimuli on each day, indicating that the longitudinally registered neurons subsequently used for analysis comprise a sufficiently representative sample of all recorded neurons. (**Supplementary Fig. 2c-j, Supplementary Table 1**). Within days, the proportion of total shock- and stable tone-and-trace responsive neurons during Extinction 1 positively correlated with freezing in early Extinction 3 in psilocybin mice, and suppressed shock-responsive neurons during Extinction 1 correlated with freezing in early Extinction 3 in saline mice, suggesting that neural response properties during Extinction 1 may be related to behavioral extinction across groups.

### Psilocybin alters dynamics and encoding of freezing behavior

The RSC was host to many neurons encoding freezing behavior in every session (**Supplementary Fig. 2b, Supplementary Table 1**). Interestingly, the average freezing encoding of individual neurons increased during *Acquisition*, suggesting that RSC neurons preferentially encode acute fear-related freezing (**Fig. 2p,q, Supplementary Table 1**). Behaviorally, psilocybin significantly reduced bout number and increased bout length acutely during Extinction 1 without affecting total freezing time, indicating an effect of psilocybin treatment on the ability to maintain sustained freezing (**Fig. 3a,b, Supp Fig. 3, Supplementary Table 1**). This could be due to an interruption of attention to or recall of fear cues.

**Figure 3.**
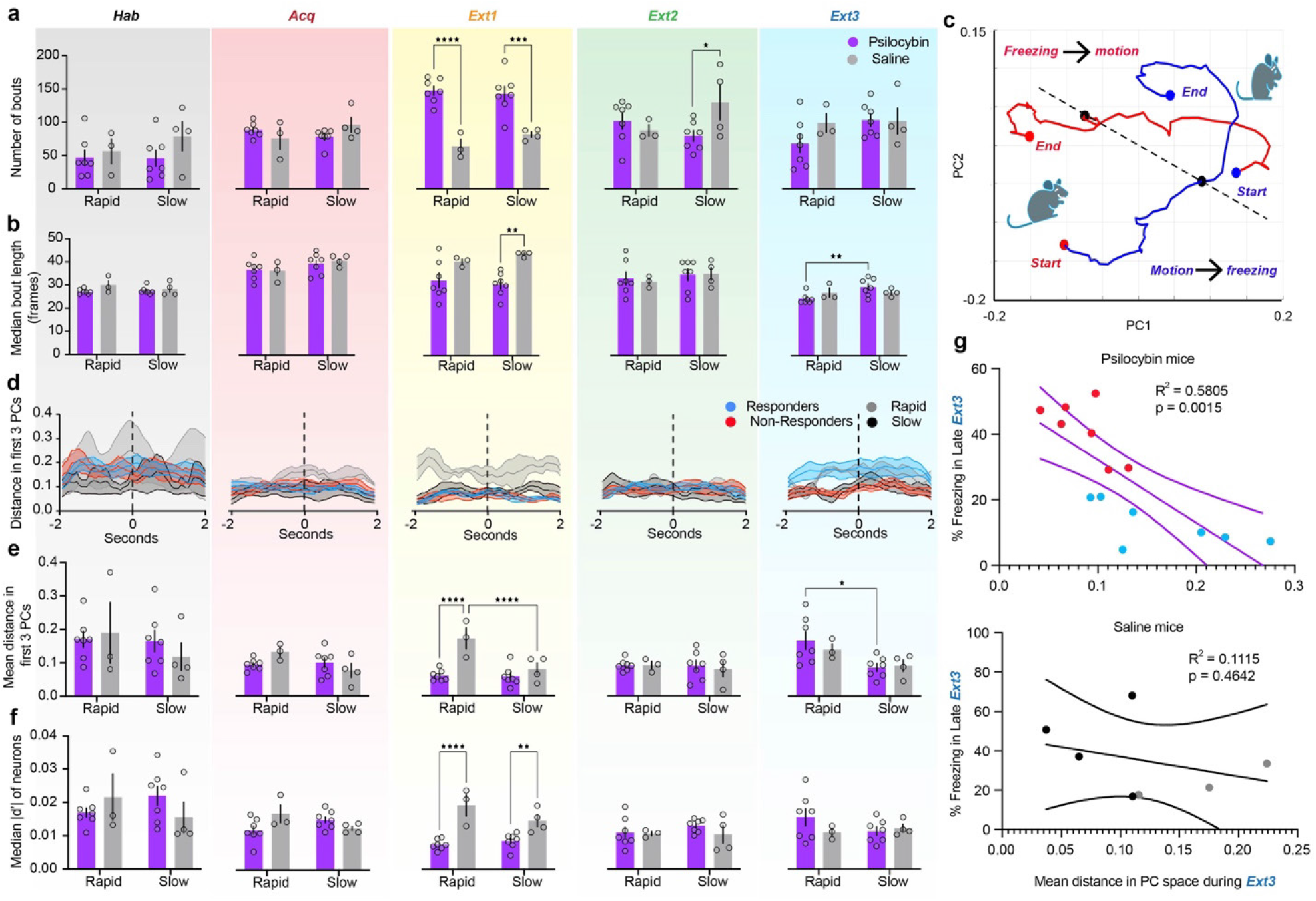
Psilocybin alters dynamics and encoding of freezing behavior. **a**. Total number of freezing bouts per session. (Left to right, Black = Habituation, Red = Acquisition, Yellow = Extinction 1, Green = Extinction 2, Blue = Extinction 3). Two-way ANOVA. (Supp. Table 1, rows 29-33). **b**. Median bout length per session in frames. Two-way ANOVA. (Supp. Table 1, rows 34-38). **c**. Representative average trajectories of motion-to-freezing (blue line, bout start) and freezing-to-motion (red line, bout end) transitions in the first two PCs, from two seconds before to two seconds after transition. (Black point = time of transition, red points = starting and ending points in motion, blue points = starting and ending points in freezing) A dashed line is drawn between the two transition points. **d**. Average Euclidean distance in PC space between each pair of points in motion-to-freezing and freezing-to-motion trajectories in the first three PCs on each day. Dashed line indicates time of behavioral transition. Shaded areas are SEM. **e**. Mean distance in PC space between bout start and bout end over the four-second time window between trajectories. Two-way ANOVA. (Supp. Table 1, rows 39-43). **f**. Median absolute value of d-prime between motion and freezing in all recorded neurons on each day. Two-way ANOVA. (Supp. Table 1, rows 44-48). **g**. Linear regression of distance in PC space between trajectories (Fig. E, right) and % trace period freezing in late Extinction 3 in psilocybin (top) and saline mice (bottom). (Supp. Table 1, rows 49-50). Data are mean ± SEM. * p ≤ 0.05, ** p < 0.01, *** p < 0.001, **** p < 0.0001.

In non-longitudinally registered neurons, transitions from freezing to motion and vice versa were robustly encoded in the first 3 PCs of RSC activity (**Fig. 3c**). Average Euclidean distance from freezing-to-motion and motion-to-freezing trajectories tended to be stable within this time window. However, distance greater during Extinction 1 in rapidly compared to slowly extinguishing saline mice, a difference occluded by psilocybin (**Fig. 3d, Supplementary Table 1**). Accordingly, single cell discriminability between freezing and motion, in terms of the population’s median magnitude of d-prime, was suppressed by psilocybin (**Fig. 3e, Supplementary Table 1**). Interestingly, population freezing was subsequently enhanced in responders during Extinction 3 and predicted freezing levels in psilocybin but not saline mice (**Fig. 3d,f, Supplementary Table 1**). Thus, while RSC discriminability between motion and freezing during Extinction 1 is enhanced in rapidly extinguishing mice, psilocybin acutely suppresses this neural discriminability preceding its rebound 48 hours later in responders. Psilocybin therefore interrupts freezing behavior and encoding in mice.

### Tensor component analysis reveals evolution of RSC through different states over fear and extinction learning

We hypothesized that psilocybin induces the rapid recruitment of a novel Extinction associated ensemble in the RSC during Extinction 1 that persistently drives RSC activity during future extinction sessions. Enhanced extinction could also arise from psilocybin-mediated suppression of specific ensembles associated with the fear learning and memory. To identify ensembles associated with TFC acquisition and extinction, we employed an unsupervised dimensionality reduction technique developed by Williams et al., 2018 called Tensor Component Analysis (TCA) that can be used to group neurons into functional ensembles defined by their within- and across-trial dynamics ^73^ (**Fig. 4a**).

**Figure 4.**
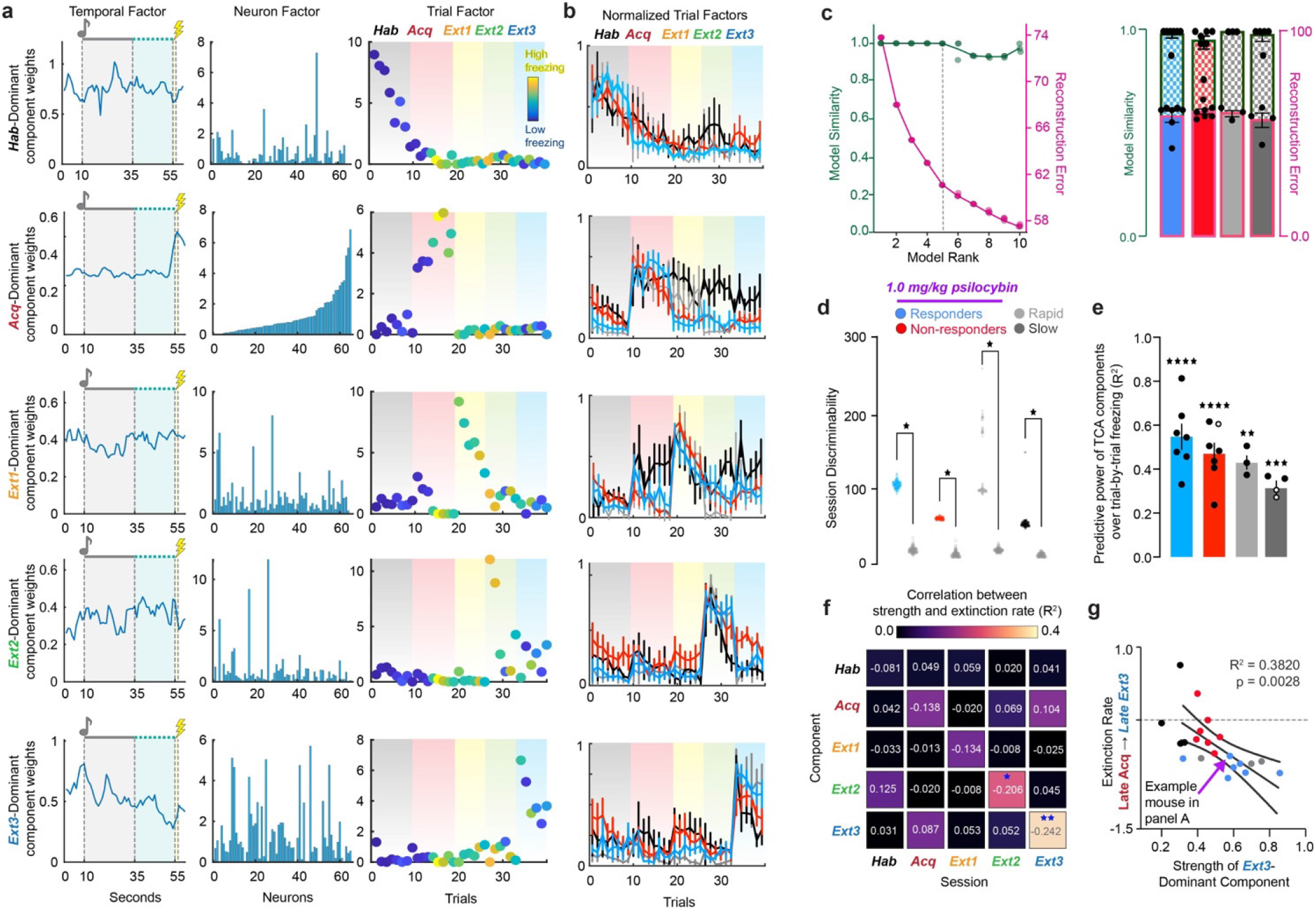
Tensor Component Analysis (TCA) captures evolution of RSC through different task-relevant states over learning. **a**. Representative rank-5 TCA model of neural activity over RSC in one mouse. Rows correspond to unique components of neural activity and columns correspond to temporal, neuron, and trial factors. Values in each panel correspond to the factor loadings, or weights, of each component at each time, cell, and trial in the given component. Pink dashed line over temporal factors indicates time of conditioned one delivery, and lightning bolt indicates time of shock-delivery. Gradients over the trial factor indicate session of trials (Black = Habituation, Red = Acquisition, Yellow = Extinction 1, Green = Extinction 2, Blue = Extinction 3). Trial weights are color coded according to the animals % time freezing in each trial (dark blue = 0%, bright yellow = 100%). **b**. Normalized trial factor weights for each component, averaged within groups. Two-Way RM ANOVA. (Supp. Table 1, rows 51-55). **c**. Validation of choice of rank-5 model. Left: Four TCA models at each of ranks 1-10 were generated on neurons pooled from all psilocybin administered animals and their reconstruction error (pink) and model similarity (blue) plotted against each other. Rank-5 was chosen by minimizing reconstruction error while maximizing model similarity (black dashed line). (Supp. Table 1, row 56) Right: Reconstruction error (solid colors) and similarity (checkered colors) in individual animals. (Supp. Table 1, row 57) Ordinary One-Way ANOVA with Tukey multiple comparisons correction. **d**. Trial weights of dominant factor during a given session divided by those of each other factor, summed over sessions, calculated over 100 iterations of TCA on real data from each group and TCA models generated on 100 shuffles of the data. Data was shuffled over cells at each individual timepoint to preserve all temporal and trial dynamics of activity that could lead to session discriminability. Multiple unpaired t-tests. P<0.0001 for all comparisons. (Supp. Table 1, row 58) **e**. Linear regression trial factor value of each of 5 components and trial-by-trial freezing across all 5 days (R^2^). Significant values are filled and non-significant values are hollow. One-sample t-test. (Supp. Table 1, rows 59-60) **f**. Linear regression of relative strength of each component during each session and extinction rate in all mice (R^2^). Numbers are linear coefficients. Stars indicate where slope is significantly non-zero. (Supp. Table 1, row 61-65) **g**. Data in D for the Extinction 3-dominant component during Extinction 3. (Supp. Table 1, rows 66) * p ≤ 0.05, ** p < 0.01, *** p < 0.001, **** p < 0.0001

To determine the appropriate model rank for analysis, TCA was run on cells pooled from all animals in each treatment group to calculate model reconstruction error and similarity as a function of increasing model rank. The elbow method revealed that models of rank 5 were most appropriate for subsequent analysis, and such models were generated for individual animals (**Fig. 4c**) Across animals, rank 5 models did not identify within-trial temporal dynamics beyond shock-responsiveness (**Supplementary Fig. 3a-e, Supplementary Table 1**). They did, however, cluster trials from the same session, identifying RSC dynamics driving distinct phases of TFC acquisition and extinction (**Fig. 4a,b**).

To eliminate the possibility that trial clustering was due to changes in recording quality between days or cell misalignment, 100 iterations of TCA on the real data from each group were compared to TCA models of 100 shuffles of the neural activity. Neural activity was shuffled by cells at each timepoint, such that the average activity over time and trials was preserved. This way, differences in between- and within-trial temporal dynamics of the entire population would be entirely conserved, but the ensembles driving those differences would be abolished.

To calculate the session discriminability of the real and shuffled TCA models, we exploited the clustering of trial factor weights within a given session, yielding a dominant component for each session. In a model of rank *R*, for component *r* in session *s* with mean trial weights 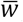, the relative strength of each component. The model’s subsequent session discriminability index were calculated as:

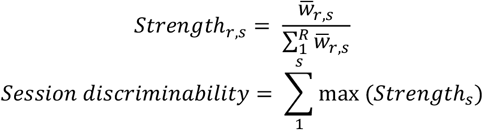

When cells were shuffled to preserve the within- and acrosstrial structure of the data, session discriminability was significantly diminished in every group, rejecting the hypothesis that same-session trial clustering was due to recording or registration artifacts (**Fig. 4d, Supplementary Table 1**). In non-shock control mice, who did not undergo any associative learning beyond neutral sensory integration and context familiarization (*i*.*e*., no electric shocks during Day 2 Acquisition), session discriminability was reduced, suggesting TCA identified task-relevant components of RSC activity **(Supplementary Fig. 5a-d, Supplementary Table 1**).

In conditioned mice, the relative strength of each component strongly also predicted freezing across and within their own session with R^2^ > 0.1 (**Fig. 4e,f, Supplementary Table 1**). The strength of the *Acquisition-*dominant component during Acquisition and Extinction 3 oppositely predicted freezing, while the *Extinction 3*dominant component during Extinction 3strongly predicted extinction rate across groups, suggesting the identification of fear extinction related neural dynamics in the RSC by TCA (**Fig. 4g, Supplementary Table 1**). These results confirm that the evolution through unique dynamics across days is a learning-related process in the RSC.

### Turnover in the dominant neural ensembles driving RSC dynamics predicts fear extinction

The neuron factor weights returned by TCA were used to identify ensembles driving the *Acquisition*-, *Extinction 1*-, and *Extinction 3*-dominant components of RSC activity in each mouse. When simulated tensors for each animal populated with identically behaving neurons, the mean and median weights were *w =* 1.0694 and 1.0709 respectively, suggesting that, if all neurons contributed equally, each neuron would be assigned *w ∼*1 (**Supplementary Fig. 6d-g**). Thus, *w=*1 was considered a reasonable null hypothesis for the strength of a neuron’s participation in each component, such that if a neuron’s weight was greater than 1, then it was included in the ensemble. 40% of neurons met this criterion for each ensemble, resulting in considerable ensemble overlap (**Fig. 5a,b, 6a, Supplementary Fig. 4a-c, Supplementary Table 1**). Ensemble overlaps are of interest as cells driving RSC dynamics on both *Acquisition* and *Extinction 1*, for instance, might in part comprise a neural substrate for a fear memory^61^.

**Figure 5.**
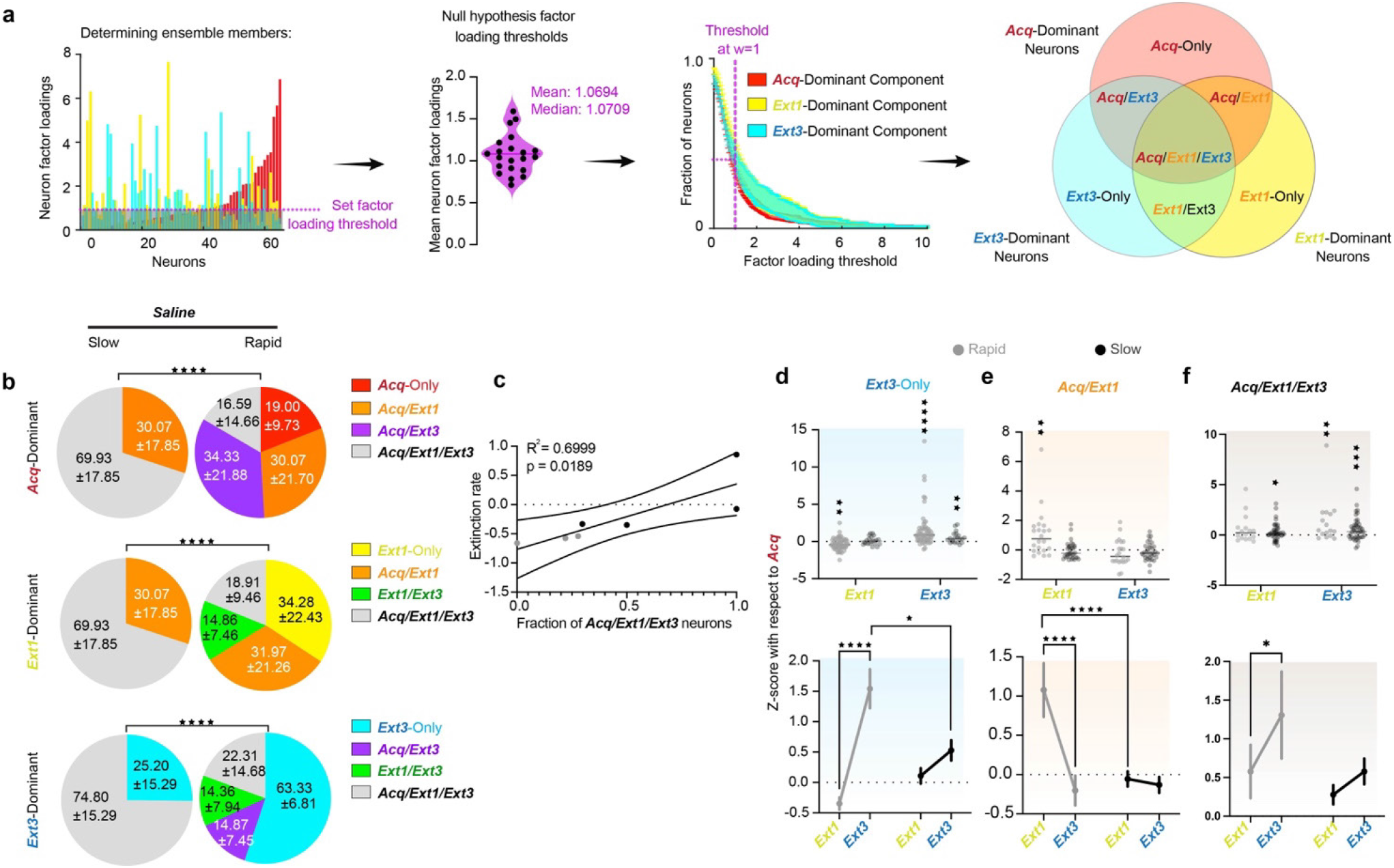
Turnover in the dominant neural ensembles driving RSC dynamics predicts fear extinction. **a**. Choosing Acq-, Ext1-, and Ext3-dominant neurons (red, yellow, and blue, respectively). Left: The fraction of neurons included in the ensemble at various thresholds across animals (mean, SEM) and the neuron factor weights of each neuron in each component in a representative animal. Neurons crossing the chosen threshold of *w=1* are indicated by enhanced opacity. Middle: Schematic of the overlaps between these neurons, yielding Acq-Only, Acq/Ext1, Ext1-Only, Ext1/Ext3, Ext3-Only, Acq/Ext3, and Acq/Ext1/Ext3. Ensembles are denoted by the corresponding ROYGBIV color code throughout the figure. Right: Example traces. **b**. Pie charts describing the average overlap of the Acq-, Ext1, and Ext3-dominant ensembles (top, middle, bottom) in rapidly and slowly extinguishing saline-administered mice. Numbers are mean ± SEM. Stars indicate comparisons between each psilocybin group and saline. Chi-square test. (Supp. Table 1, rows 67-69). **c**. Linear regression of the fraction of *Acq/Ext1/Ext3* neurons and extinction rate in saline mice. (Supp. Table 1, row 70). **d**. Top: z-score activity in individual *Ext3-only* neurons in each ensemble from Acquisition. Wilcoxon rank-sum to test if change is different from zero. Bottom: Same data displayed as mean ± SEM. Two-way RM ANOVA to compare changes over time and between groups. (Supp. Table 1, rows 71-72). **e,f**. Same as D for *Acq/Ext1* and *Acq/Ext1/Ext3*. (Supp Table 1, rows 73-76). * p ≤ 0.05, ** p < 0.01, *** p < 0.001, **** p < 0.0001.

**Figure 6.**
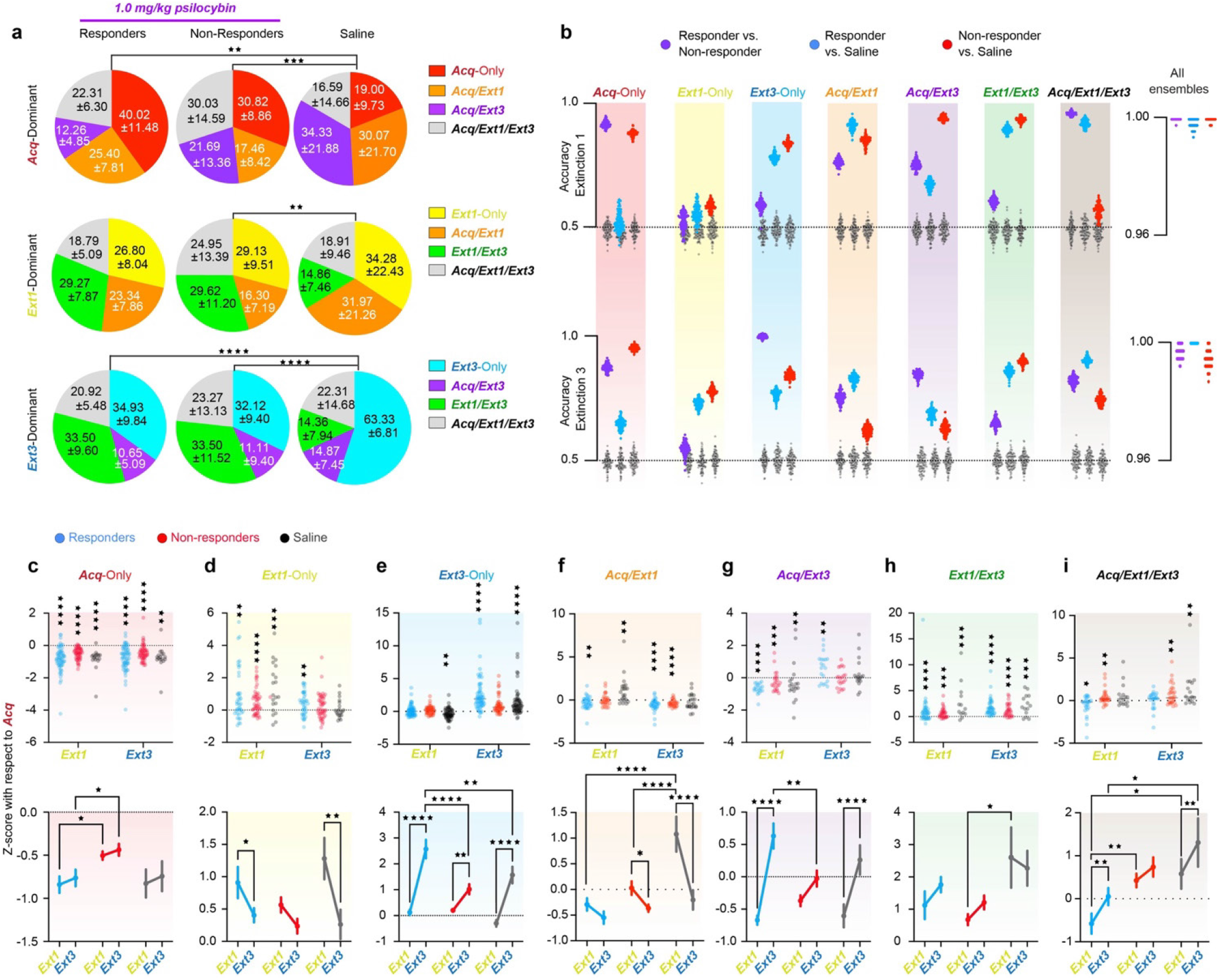
Psilocybin bidirectionally modulates neural ensembles driving RSC dynamics during TFC in responders. **a.** Pie charts describing the average overlap of the Acq-, Ext1, and Ext3-dominant ensembles (top, middle, bottom) in responders, non-responders and rapidly extinguishing saline-administered mice. Numbers are mean ± SEM. Stars indicate comparisons between each psilocybin group and saline. Chi-square test. (Supp. Table 1, rows 77-79) **b.** Accuracies of 100 Fisher decoders trained to predict responder status (left cloud, purple), responders from rapidly extinguishing saline-administered mice (middle cloud, blue around grey), and non-responders from saline administered mice (right cloud, red around grey). Grey clouds are the same decoders tested on shuf?ed class labels. Decoders were trained on activity during Extinction 1 (top) and Extinction 3 (bottom). Right-hand panels accuracies of decoders trained on all seven ensembles as predictors. **c.** Top: z-score activity in individual Acq-Only neurons in each ensemble from Acquisition. Wilcoxon rank-sum to test if change is different from zero. Bottom: Same data displayed as mean ± SEM. Two-way RM ANOVA to compare changes over time and between groups. (Supp. Table 1, rows 80-81) **d-i.** Same as C for Ext1-Only, Ext3-Only, Acq/Ext1, Ext1/Ext3, Acq/Ext3, and Acq/Ext1/Ext3, respectively. (Supp. Table 1, rows 82-93) * p ≤ 0.05, ** p < 0.01, *** p < 0.001, **** p < 0.0001

**Figure 6.**
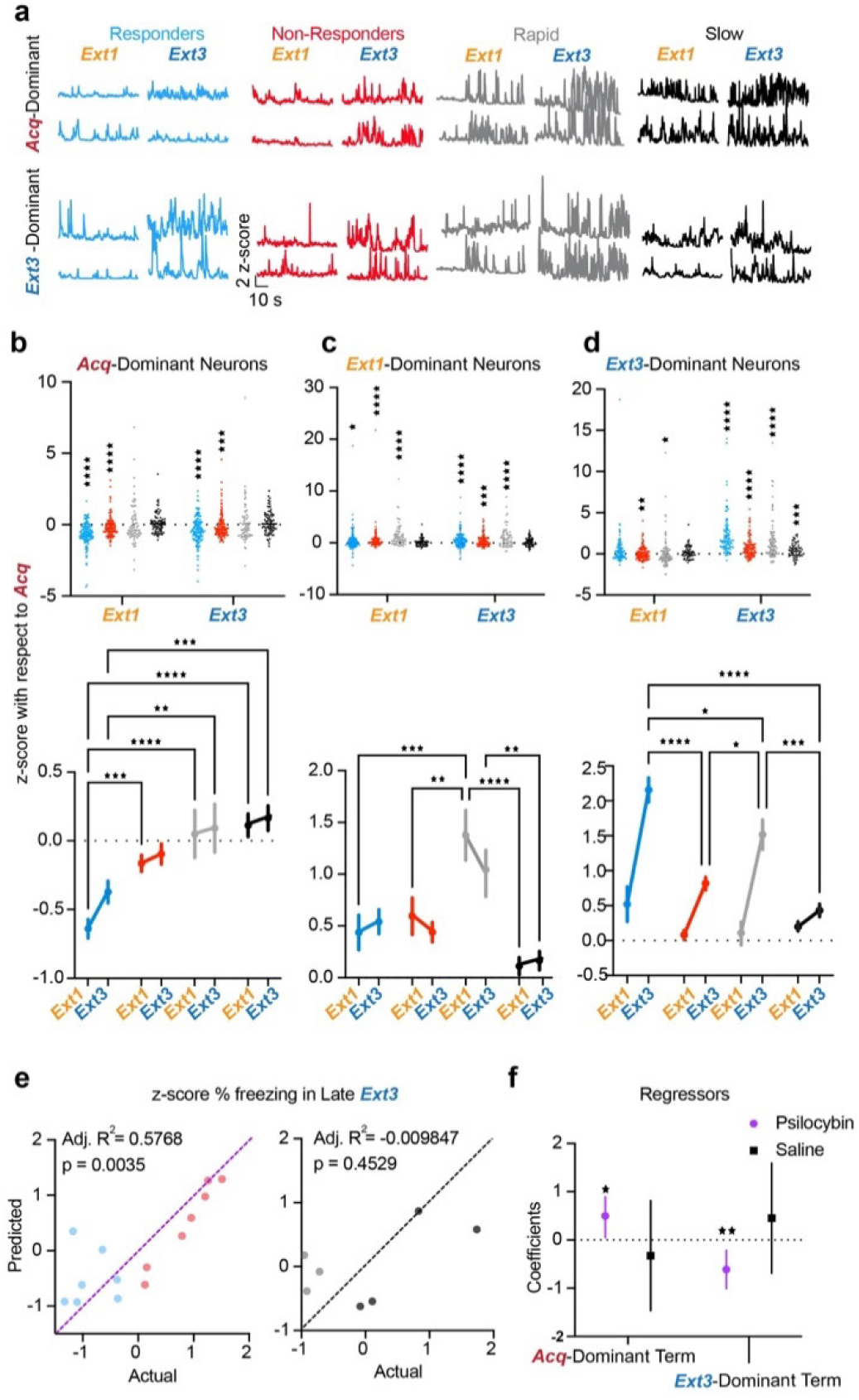
Psilocybin induces long-term suppression of Acqdominant neurons and strong post-acute recruitment of Ext3dominant neurons in responders. **a**. Example traces of Acq-dominant (top) and Ext3-dominant (bottom) neurons during Extinction 1, and Extinction 3 in each group. **b**. Top: z-score with respect to Acquisition of individual Acq-dominant neurons in each ensemble during Extinction 1 and 3. Wilcoxon rank-sum to test if median ≠ 0. Bottom: Same data displayed as mean ± SEM. Two-way RM ANOVA to compare changes over time and between groups. Data are represented as mean ± SEM. (Supp. Table 1, rows 94-95) **c**. Same as B) for Ext1-dominant neurons. (Supp. Table 1, rows 96-97). **d**. Same as B) for Ext3-dominant neurons. (Supp. Table 1, rows 98-99) **e**. Multiple regression of z-score % freezing in late Extinction 3 on z-score from Acquisition of activity of *Acq-*dominant neurons in Extinction 1 and *Ext3*-dominant neurons in Extinction 3 in psilocybin (left) and saline mice (right). (Supp. Table 1, rows 100-1) **f**. Mean ± 95% confidence intervals of regression coefficients. (Supp. Table 1, row 102) * p ≤ 0.05, ** p < 0.01, *** p < 0.001, **** p < 0.0001

Intriguingly, although every combination of these ensembles (*Acq-Only, Ext1-Only, Ext3-Only, Acq/Ext1, Acq/Ext3, Ext1/Ext3, and Acq/Ext1/Ext3*) were represented rapid saline mice, slow saline mice failed to recruit any new dominant neurons in Extinction 1, resulting in a large *Acq/Ext1/Ext3* ensemble and smaller *Acq/Ext1* and *Ext3-only* ensembles (**Fig. 5b, Supplementary Table 1**). Rapidly extinguishing mice recruited a new Extinction 1dominant ensemble and suppressed a portion of the Acquisition dominant ensemble (19%, *Acq-only* neurons). However, slowly extinguishing mice did neither until Extinction 3, where only 25% of neurons were recruited compared to 55% in rapidly extinguishing mice. The proportion of *Acq/Ext1/Ext3* neurons strongly predicted extinction rate in saline mice, confirming the interpretation that ensemble turnover is associated with effective fear extinction (**Fig. 5c, Supplementary Table 1**).

Tracking the change in activity in these ensembles from Acquisition revealed that slowly extinguishing mice display blunted plasticity in the *Ext3-*only and *Acq/Ext1* neurons compared to rapidly extinguishing mice (**Fig. 5d,e, Supplementary Table 1**). While *Acq/Ext1/Ext3* neurons are significantly more activated during Extinction 1 and Extinction 3, rapid mice display a strong enhancement of *Acq/Ext1* and *Ext3-only* activity during Extinction 1 and 3 respectively, while *Acq/Ext1* neurons are maintained, and *Ext3-only* neurons weakly enhanced in slowly extinguishing mice. Instead, an enhancement *Acq/Ext1/Ext3* activity appears to be the defining feature of RSC activity in slow-extinguishing mice (**Fig. 5f, Supplementary Table 1**). Thus, both in terms of neuronal identity and activation, plasticity in dominant RSC ensembles appears to be a feature of effective TFC extinction.

### Psilocybin enhances extinction-associated ensemble turnover

Both responders and non-responders exhibited high ensemble turnover, in contrast to slowly extinguishing saline mice. To investigate the effect of psilocybin on plasticity of RSC ensembles, responders and non-responders are compared only to rapid saline mice, due to the nonexistence of these ensembles in slowly extinguishing mice. Ensemble overlaps significantly differed between psilocybin and rapid saline mice in most cases, but not between responders and non-responders (**Fig. 6a top, Supplementary Table 1**). The overlap between *Acq-* and *Ext1-*dominant neurons was similar across all groups. Non-responders exhibited the greatest proportion of *Acq/Ext1/Ext3* neurons, while rapid saline mice exhibited the greatest proportions of *Acq/Ext1* and *Acq/Ext3* neurons. Responders displayed the greatest proportion of *Acq-only* neurons, the only ensemble defined by its persistent suppression during both Extinction 1 and Extinction 3. Though only a greater proportion of the *Ext1-*dominant ensemble is defined by newly recruited neurons in psilocybin mice than in rapid saline mice, (**Fig. 6a middle**). Psilocybin mice recruited double proportion of *Ext1/Ext3* neurons. Finally, while more neurons were newly recruited in the *Ext3-*dominant ensemble in rapid saline mice, similar proportions of neurons had been recruited after Acquisition (**Fig. 6a bottom, Supplementary Table 1**). The proportion of *Ext1/Ext3* neurons comprising the Extinction 3-dominant ensemble was triple that of saline mice in psilocybin mice. Thus, psilocybin acutely accelerates a rapid turnover from fear Acquisition-dominating neurons to a stable Extinction-dominant population. This result is highly consistent with the hypothesis that psilocybin both establishes and stabilizes novel neural ensembles. In non-shock controls, the proportion of *Acq*/*Ext1* and *Acq/Ext1/Ext3* neurons was much smaller than in saline mice, while there were more *Ext1/Ext3* neurons. (**Supplementary Fig. 5e-g, Supplementary Table 1**). These result supports the hypothesis that preferential overlaps of *Acq/Ext1-* and *Ext1/Ext3*-dominant ensembles, in saline and psilocybin mice respectively, indicate enhanced stability of fear acquisition- and extinction-related ensembles over time.

### Activity in neural ensembles predicts treatment and responder status

Fisher linear decoders were trained to distinguish between psilocybin responders, non-responders, or saline mice based on the average activities of each identified ensemble during either *Extinction 1* or *Extinction 3* (**Fig. 6b, Supplementary Fig. 4d-e**). Decoders were trained to classify two groups at a time – responders vs. non-responders, responders vs. saline, non-responders vs. saline to determine which ensembles varied with treatment, extinction class, or both.

During Extinction 1, when psilocybin mice were under acute influence of the drug, the *Ext3-Only* and *Ext1/Ext3* ensembles specifically distinguished both groups of psilocybin mice from saline mice, suggesting that psilocybin affected activity in these ensembles in a behavior-nonspecific fashion (**Fig. 6b top**). The *Acq/Ext1* and *Acq/Ext3* ensembles discriminated between all groups, suggesting that psilocybin’s acute effects on these neurons can predict future behavioral change. The *Acq/Ext1/Ext3* ensemble only dis criminated between responders and the other two groups, suggesting that, while this ensemble is not determinately affected by psilocybin, altered activity in this ensemble during psilocybin administration may enhance future behavioral change. The *Acq-Only* ensemble significantly distinguished nonresponders from the other two groups, suggesting that altered activity in this ensemble under psilocybin may block future behavioral change. During *Extinction 3*, the *Ext3-Only* ensemble came to distinguish all three groups (**Fig. 6b bottom**). These results suggest that psilocybin acutely alters dynamics in these neurons acutely and post-acutely in a manner predicting behavior.

When models were trained on all seven ensembles as predictors, they predicted treatment and responder status with > 95% accuracy on all 100 iterations for each pair (**Fig. 6b right**). Classification between all three groups verified these results (**Supplementary Fig. 4e-f**). The ability of many ensembles to distinguish responder status during Extinction 1 suggests that neural activity in the RSC during psilocybin exposure may be a crucial determinant of therapeutic-like response 48 hours later.

### Acute suppression of *Acq-*dominant neurons and delayed recruitment of *Ext3-*dominant neurons predict psilocybin-enhanced extinction

To explore the development of the distinctive predictive characteristics of each ensemble, we calculated how the activity of each neuron in these ensembles changed from the Acquisition session. *Acq-Only* neurons were suppressed during Extinction 1 and Extinction 3 in all groups, but significantly less so in non-responders, suggesting that the suppression of this ensemble during early extinction may affect the pace of fear extinction in mice (**Fig. 6c, Supplementary Table 1**). *Ext1-Only* neurons were potentiated in all groups during Extinction 1, but only remained significantly greater than zero during Extinction 3 in responders (**Fig. 6d, Supplementary Table 1**).

However, this ensemble had limited predictive abilities regarding both responsiveness and treatment, weakening the claim that this difference is crucial for psilocybin’s effects on TFC extinction. *Ext3-Only* neurons were strongly recruited in all groups in Extinction 3, but significantly more greatly in responders, suggesting that this enhanced activation of *Ext3-Only* neurons 48 hours after drug administration may drive enhanced TFC extinction in responders (**Fig. 6e, Supplementary Table 1**).

*Acq/Ext1* neurons were significantly suppressed during Extinction 1 in responders, unchanged in non-responders, and enhanced in rapidly extinguishing mice, underlying this ensembles’ ability to distinguish between all three groups (**Fig. 6f, Supplementary Table 1**). This result suggests that, although psilocybin results in the suppression of *Acq/Ext1* neurons 48 hours after drug administration in both responders and non-responders, it may only ultimately enhance extinction when *Acq/Ext1-*dominant neurons are suppressed during acute drug effects. The *Acq/Ext3*dominant ensemble was significantly suppressed in Extinction 1 in all animals, and subsequently strongly potentiated with respect to Acquisition levels in Extinction 3 in responders and rapid saline mice, suggesting that these neurons were suppressed during acute drug effects and subsequently re-recruited in responders (**Fig. 6g, Supplementary Table 1**). Likewise, the *Ext1/Ext3*-dominant ensemble was potentiated across days with respect to Acquisition in all groups, but most greatly in rapid saline mice, potentially compensating for its reduced numbers in saline mice (**Fig. 6h, Supplementary Table 1**). Finally, the *Acq/Ext1/Ext3* ensemble driving activity in all three sessions was suppressed during Extinction 1 in responders but potentiated in non-responders and saline mice (**Fig. 6i, Supplementary Table 1**). This result suggests that acute suppression of this ensemble during psilocybin administration may enhance the likelihood of enhanced TFC extinction.

For a holistic picture of these results, one can consider the entire *Acq*-, *Ext1*-, and *Ext3-*dominant ensembles. In saline mice, the *Acq-*dominant ensemble is persistently active at Acquisition-like levels throughout Extinction, regardless of extinction rate (**Fig. 7b, Supplementary Table 1**). This result is specific to TFC-trained mice, as opposed to non-shock mice, indicating the persistence of a potential substrate for fear memory throughout extinction in saline mice (**Supplementary Fig. 5h, Supplementary Table 1**). In contrast, psilocybin persistently suppresses the *Acq-*dominant ensemble, strongly in responders and weakly in non-responders (**Fig. 7b, Supplementary Table 1)**. The *Ext1-*dominant ensemble is potentiated throughout extinction in all groups, though most in rapid mice due to the lack of inhibition of overlapping *Acq-*dominant neurons, unlike psilocybin mice, and the presence of newly recruited neurons, unlike slowly extinguishing mice (**Fig. 7c, Supplementary Table 1**). Finally, recruitment of the *Ext3-*dominant ensemble more greatly in saline mice than non-shock controls, suggesting that heightened activity in novel RSC ensembles is a feature of TFC extinction (**Fig. 7d, Supplementary Table 1**). However, the *Ext3*-dominant ensemble is most greatly recruited in both responders and rapidly extinguishing saline mice, suggesting that this recruitment is a fixed feature of effective fear extinction in the RSC, regardless of treatment. However, its recruitment was enhanced in psilocybin responders compared to rapid saline mice. Critically, these results were robust to varying the factor loading thresholds determining a neuron’s ensemble membership (**Supplementary Fig. 6a-c, h**).

To determine the relationship between changes in neural activity and behavior, we regressed percent time freezing in the last half of Extinction 3 on the change in neural activity of the *Acq-* dominant ensemble during Extinction 1 and the *Ext3*-dominant ensemble of Extinction 3. (**Fig. 7e,f**) We found that these variables both significantly and oppositely predicted freezing in psilocybin mice but not saline mice. Thus, psilocybin may enhance TFC extinction in animals by bidirectionally modulating ensembles underlying distinct phases of TFC.

### A computational model of a two-ensemble RSC microcircuit explains psilocybin’s effects

*Acq-*dominant neurons are largely shock-responsive (**Fig. 8a**). Ideally, to test their causal influence on psilocybin-enhanced extinction, we would capture and manipulate this functional ensemble, with methods such as TRAP or scFlare paired with optoor chemogenetics. These approaches require that the targeted neurons have largely homogenous and stable response properties, distinct from the general population, across trials. However, we found that most neurons significantly respond to the shock on only 1-3 of the total 8 trials (**Supplementary Fig. 7a,d**). Importantly, on each trial, the set of shock-responsive neurons only overlaps by 30-40% with any other trial (**Supplementary Fig. 7b**), while a similar proportion of the general population are also shock responsive (**Supplementary Fig. 7c**). Thus, this functional population does not meet the criteria necessary for these techniques, as TRAP or scFlare tagging would be nonspecific and insensitive to *Acq-*dominant neurons with trial-varying response properties.

**Figure 8.**
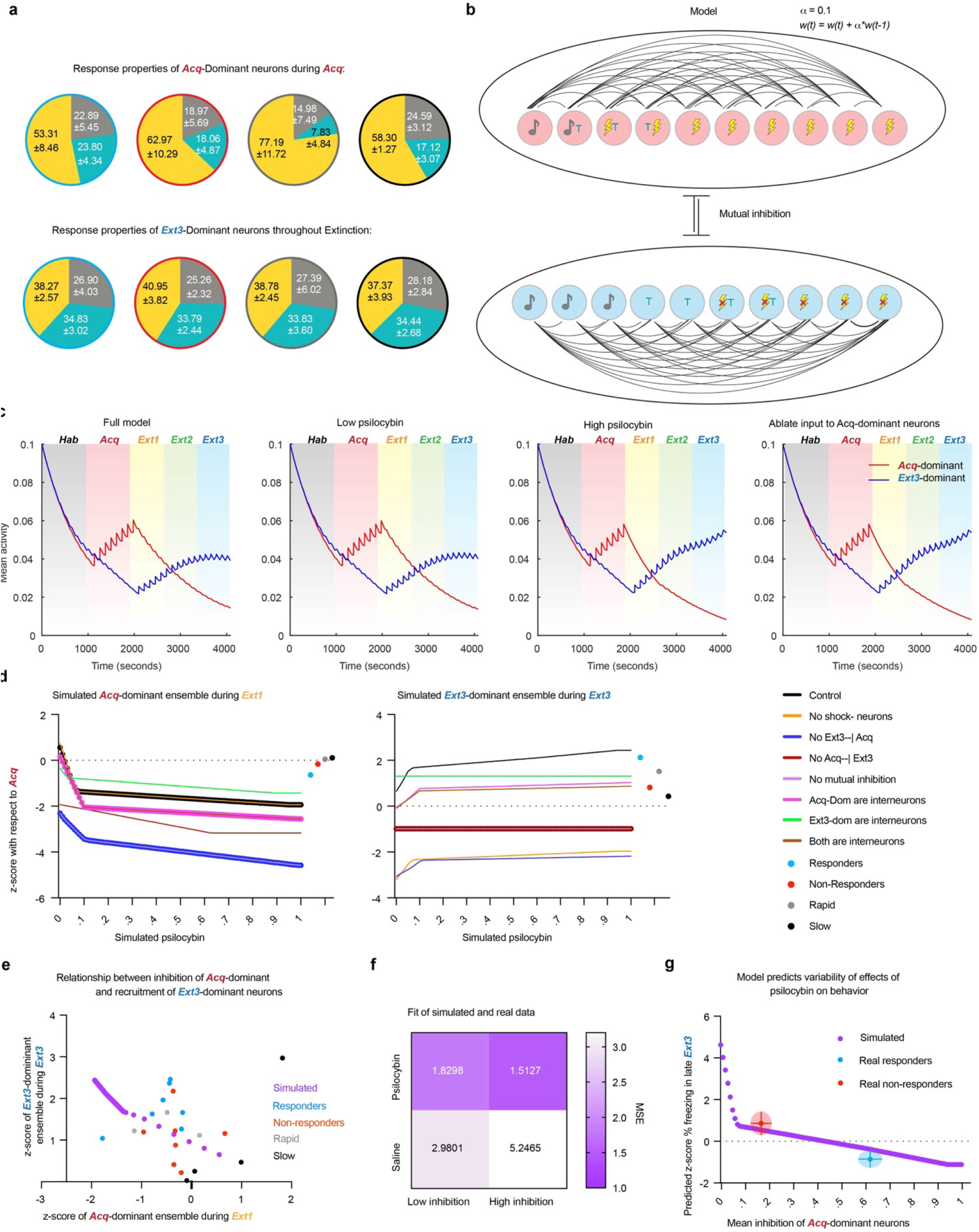
A computational model of a two-ensemble RSC microcircuit explains psilocybin’s effects. **a**. Fractions of tone, trace, and shock responsive neurons in the *Acq-*dominant ensemble during Acquisition (top) and *Ext3*-dominant neurons over all extinction sessions. **b**. Diagram of computational model of *Acq-* and *Ext3*-dominant ensembles and learning function. **c**. Average activity of ensembles over each simulation of the full model with no simulated psilocybin (first panel), low psilocybin simulated as direct synaptic inhibition to *Acq*-dominant units (second panel) during Extinction 1, high psilocybin during Extinction 1 (third panel), and with all synaptic input to the *Acq*-dominant units ablated during during Extinction 1 (last panel). **d**. Z-score with respect to Acquisition of *Acq-*dominant neurons during Extinction 1 (left) and of *Ext3-*dominant neurons during Extinction 3 (right) over multiple conditions: the full model (black), without shock-omission sensitive neurons (yellow), without inhibition of *Acq-*dominant units by *Ext3*-dominant units (deep blue), without inhibition of *Ext3-*dominant units by *Acq*-dominant units (deep red), without mutual inhibition (purple), if *Acq*-dominant units were inhibitory interneurons (pink), if *Ext3*-dominant units were inhibitory interneurons (green), if all units were inhibitory interneurons (brown). Mean values of real data are plotted to the right of each plot. **e**. Real and simulated z-score with respect to Acquisition of *Ext3-*dominant neurons during Extinction 3 plotted as a function of *Acq-*dominant neurons during Extinction 1. Simulated data (purple), responders (blue), non-responders (red), rapid mice (gray), and slow mice (black). **f**. Mean squared error (MSE) of the real data (rows) from the simulated data (columns) in E. **g**. Simulated z-score freezing late Ext3 in model as a function of amount of synaptic inhibition (purple). Average freezing and average inhibition of *Acq-*dominant neurons in responders (blue) and non-responders (red). Error bars and clouds are SD.

Therefore, to determine whether inhibition of the *Acq-*dominant ensemble by psilocybin could be sufficient to explain either enhanced recruitment of the *Ext3*-dominant ensemble and/or behavioral variability in extinction, we built a linear-nonlinear firing rate model of a hypothesized RSC microcircuit. Each of the two ensembles was modeled with ten recurrently connected units undergoing Hebbian plasticity, with heterogeneous response properties mimicking the real data. These included eight shock responsive units in the *Acq-*dominant ensemble and five shock omission-responsive units in the *Ext3*-dominant ensemble (**Fig. 8a,b**). Crucially, each ensemble inhibited the other. Psilocybin was simulated as varying amounts of direct synaptic inhibition ranging between 0 and 1.

The average activity of the *Acq*-dominant units spiked during each shock-delivery and increased over Acquisition (**Fig. 8c**). Under no and low amounts of psilocybin inhibition, *Acq*-dominant units also spiked during the first trial of Extinction 1 before gradually reducing their activity over time, demonstrating successful modeling of fear conditioning (**Fig. 8c, panels 1 & 2**). Subsequently, the shock-omission-sensitive *Ext3*-dominant ensemble elevated its activity over Extinction. This result demonstrates that the recruitment of *Ext3*-dominant neurons over extinction is a feature of this mutually inhibitory circuit, as our data suggest.

At the highest dose of inhibition and in the positive control case of entirely ablating inputs to the *Acq*-dominant ensemble, the conditioned response of the *Acq*-dominant ensemble during Extinction 1 is eliminated and activity of the *Ext3*-dominant ensemble is enhanced in Extinction 3 (**Fig. 8c, panels 3 & 4**). Therefore, inhibition of *Acq-*dominant ensembles is sufficient to explain enhanced recruitment of *Ext3-*dominant neurons.

To determine whether the assumptions in this model are necessary for its validity, we assessed the activity changes of the ensemble while systematically varying underlying assumptions about the circuit’s architecture (**Fig. 8d**). Importantly, for the model to be valid, the case of no inhibition must account for saline mice who undergo recruitment of *Ext3-*dominant ensemble without inhibition of the *Acq*-dominant. This criterion rules out every case other than our full model, where *Acq*and *Ext3*-dominant ensembles are defined as excitatory populations that mutually inhibit one another.

To determine how well our full model explains our results, we plotted the activity of the *Ext3-*dominant ensemble during Extinction 3 as a function of that of the *Acq-*dominant ensemble during Extinction 1 alongside the average values of each mouse (**Fig. 8e**). The model predicts that there is a nonlinear relationship between inhibition of *Acq-*dominant neurons and recruitment of *Ext3-*dominant neurons, comprised of a shallow quasi-linear part ranging over lower inhibition and a steep quasi-linear part ranging over greater inhibition. To determine which part fits psilocybin or saline data, we calculated the slopes of these lines and the mean squared error (MSE) to our real data (**Fig. 8f**) We found that while both parts strongly fit the psilocybin data, they more weakly fit the saline data, as expected. Compellingly, while the high inhibition part fit the psilocybin data best (MSE_high inhibition_ = 1.5127, MSE_low inhibition_ = 1.8298), the low inhibition part fit the saline mice better (MSE_high inhibition_ = 5.2465, MSE_low inhibition_ = 2.9801). These results are consistent with our observations that 1) inhibition of *Acq-*dominant ensembles is only partially present in rapid mice, while it is a consistent phenomenon in psilocybin mice (**Fig. 6c-i**) and 2) degree of inhibition of *Acq*-dominant neurons is only related to degree of subsequent recruitment of *Ext3-*dominant neurons in psilocybin mice (**Fig. 7b-d**).

Finally, to determine whether the model explains the distribution of behavioral variability of freezing in late Extinction 3, we calculated the simulated z-score % freezing using the multiple regression model from Fig. 7e,f (**Fig. 8g**). Indeed, we found that the steep, high inhibition part of the model explains the reduction of freezing in responders as a function of mean inhibition of the *Acq-* dominant ensemble and the failure do so in non-responders, with the predicted freezing lying within a standard deviation of the mean values for each group.

## Discussion

In this study, we combined *in-*vivo single cell calcium imaging of cortical ensembles with behavioral pharmacology to elucidate the neural correlates of psilocybin-enhanced extinction. The existence psilocybin-responsive and non-responsive subpopulations of humans and rats has been the subject of recent investigation^28,75^. Here, we report for the first time that mice are divided into psilocybin responsive and non-responsive groups with respect to post-acute enhancement of TFC extinction. In drug-responsive animals, psilocybin enhances expression of extinction 24 and 48 hours later. In non-responsive animals, psilocybin has no effect on behavior compared to extinction rate-matched saline animals. Acutely, psilocybin increased freezing bouts while decreasing freezing length, suggesting an acute disruption of attention or recall processes. In miniscope-implanted mice, these behavioral changes were accompanied by a reduction of single cell and population-level discriminability between freezing and motion, raising the intriguing possibility that psilocybin could acutely impair the perception of self-motion. Freezing-encoding then recovered in Extinction 3 and predicted freezing in psilocybin mice, suggesting that recovery of freezing encoding is a biomarker of effective psilocybin-modulated fear extinction.

We used TCA to identify trial-varying components of neural activity associated with fear extinction and putative task-relevant ensembles^73^. Consistent with the hypothesis that the RSC preferentially encodes the cognitive or behavioral context associated as opposed to explicit sensory events, the trial factor weights of these components tended to cluster trials from the same session without clear organization of the temporal factor weights across animals, a characteristic significantly reduced in non-shock controls. As the RSC is also involved in contextual fear conditioning and may therefore exhibit similar neural correlates in both paired and unpaired conditioning protocols, non-shock controls isolate fear acquisition- and extinction-related signals from those associated with neutral contextual novelty, exploration, and integration^39-71^.

The RSC generates, hosts, and updates engrams for consolidated fear and extinction memory. For instance, freezing can be evoked in a novel context by optogenetically reactivating RSC-tagged neurons that were initially active during fear learning in another context (*i*.*e*., akin to the Acquisition session responsive cells)^51,56^. In the present study, candidate neurons for engrams related to fear recall and extinction should be comprised of overlaps between the *Acq-, Ext1-*, and *Ext3-*dominant ensembles, such that a subset of *Acq/Ext1* and *Acq/Ext1/Ext3* ensembles participates in a fear engram, while a subset of *Ext1/Ext3* or *Ext3-only* neurons comprise an extinction engram. The behavioral relevance of these ensembles is demonstrated by accurate decoding between responders, non-responders, and rapid saline-treated mice based on each ensemble’s activity.

Consistent with previous findings and as the RSC is necessary for TFC, we found a high proportion of overlapping *Acq*/*Ext1* and *Acq/Ext1/Ext3* neurons in saline mice^61^. Intriguingly, psilocybin reduced this proportion, while the dominant ensembles in slow saline mice were solely comprised of these neurons. The predominance of these ensembles at the expense of *Acq-only, Ext1only*, and *Ext1/Ext3* neurons in slowly extinguishing saline mice, and the predictive power of the proportion of *Acq/Ext1/Ext3* neurons over extinction rate in saline mice, support the interpretation that the maintenance of fear experience-dominating neurons maintains fear-related behavior. The reduction in putative “fear memory” neurons in psilocybin mice resulted in a doubling of *Acq-Only* neurons, suggesting that psilocybin induced a robust turnover in the composition of the ensembles driving RSC activity in the Extinction 1 session. Finally, a greater proportion of *Ext1/Ext3* neurons were observed in psilocybin than saline mice, suggesting that the ensembles recruited under psilocybin were more stable in the days to come. This observation agrees with dendritic and synaptic plasticity studies that demonstrate psilocybin rapidly induces the formation and subsequent long-term stabilization of behaviorally relevant neural pathways. However, the lack of association between size of these ensembles with extinction rate ruled out the possibility that ensemble turnover alone influences psilocybin responsiveness.

Previous work showed that that novel ensembles are recruited in the RSC during fear extinction^56^. Indeed, we observed a substantial recruitment of neurons unique to the *Ext3*-dominant ensemble in all groups, with significantly lesser activation in nonshock mice. These neurons were significantly more strongly activated during Extinction 3 in psilocybin responders than in any other group. However, this recruitment was preceded by the acute and robust inhibition of *Acq*-dominant neurons in psilocybin responders that was strikingly absent in saline mice. We found that both the overall inhibition of the *Acq-*dominant ensemble during psilocybin administration and the subsequent recruitment of *Ext3-*dominant neurons during Extinction 3 strongly predicted freezing in late Extinction 3 in psilocybin mice, but not saline mice. Furthermore, by modeling these ensembles as mutually inhibitory populations comprised of excitatory neurons, we not only replicated the neural dynamics observed in saline mice but demonstrated that varying the acute inhibition of *Acq-*dominant neurons during Extinction 1 is sufficient for enhanced recruitment of the *Ext3-*ensemble and extinction in principle.

All other versions of the model where the underlying assumptions of circuit architecture were altered failed to explain the recruitment of *Ext3*-dominant neurons in the absence of *Acq-*dominant inhibition observed in saline mice. Therefore, we can conclude that 1) *Acq-* and *Ext3-*dominant ensembles are likely polysynaptically, mutually inhibitory excitatory populations in the RSC and 2) the inhibition of *Acq-*dominant neurons is sufficient to explain neural and behavioral variability in our task in response to psilocybin. Given all mice received the same dose of psilocybin, these results raise the exciting possibility that individual differences in receptor availability or circuit anatomy in the RSC facilitate psilocybin-enhanced fear extinction.

These results complicate the prevailing hypotheses in the field that psilocybin’s effects on behavioral flexibility are downstream of excitatory activity and plasticity enacted via 5HT2ARs^8^. To the contrary, we observed no increased activity under acute psilocybin. As the RSC is rich in 5HT2CRs and 5HT1Ars, both inhibitory with high binding affinity to psilocin, it is plausible that psilocin directly inhibits neurons in this region^28,60,61,76.^ Alternatively, psilocin could excite inhibitory neurons upstream of *Acq-* dominant neurons to exert its effects. Therefore, it is possible that behavioral variability under psilocybin in this task could be due to variability in receptor expression and availability or in anatomical connectivity.

Identifying these neurons will be key to establishing their causal influence on fear extinction with or without psilocybin and potentially developing targeted therapeutics. One possibility is that these neurons are salience or valence-sensitive. Notably, a single-dose of psilocin reduces neural activities to aversive airpuffs in the central amygdala days later, hinting that weakened neural activities within negative valence-encoding circuits may partially contribute to this observation^77^. In the RSC, our data and model suggest that *Acq-* and *Ext3-*dominant ensembles contain neurons responsive to shock or shock omission, stimuli of opposite valence. Furthermore, our observations echo the finding that inhibitory plasticity in hippocampus fear memory engrams is necessary for the development memory selectivity, measured by the reduction of freezing in neutral contexts over time^78^. As RSC neurons can encode changes in reward values^47,48^, ensembles in this study could constitute a valence- or salience-sensitive ensembles in the RSC, a feature that can enable their genetic and anatomical identification. Taken together, these results suggest that psilocybin both enhances endogenous mechanisms of fear extinction – the potentiation of newly recruited RSC neurons – and, or possibly because, it engages non-typical mechanisms as well – the suppression of fear acquisition-dominating neurons in drug responders. These results support a current field hypothesis that the neurophysiological effects of psychedelics underlying behavioral flexibility involve altering task-relevant activity in neural ensembles over subsequent days^79^. However, rather than simply accelerating or enhancing endogenous mechanisms of behavioral flexibility (*i*.*e*., increasing activity in new ensembles), psilocybin also engages an inhibitory mechanism of fear extinction. Indeed, the acute, response-predicting effects of psilocybin observed in this study are entirely comprised of inhibition of fear acquisition-associated neurons. Psilocybin’s enhancement of extinction-like activity is not observed until the days following treatment and can be explained by prior suppression of fear memory-associated activity. Future research will explore how the neuroplastic effects of psilocybin on a cellular and circuit level evoke these distinct effects on neural dynamics and establish a causal relationship between the ensemble-specific changes in activity observed here with behavior.

## Methods

### Experimental Methods

#### Animals

Animals used in all studies were C57BL/6J mice *Animals:* Animals used in all studies were C57BL/6J mice from Jackson Laboratories (RRID: IMSR_JAX:000664). Mice were kept on a reverse 12-hour light/dark cycle. Behavior was performed at least 1 hour and no more than 4 hours following lights-off. Group-housed males (n=34) and females (n=16) between 8-12 weeks of age were used in the behavioral pharmacology experiment in Fig. 1. For the Miniscope study, males of 8-10 weeks of age underwent viral injection surgeries, followed by implantation of 4.0 (length) x 1.0mm (diameter) GRIN lens at 10-12 weeks, and behavior at 12-16 weeks (minimum 2-week recovery time from last surgery). Mice were singly housed following implant surgery.

#### TFC Conditioning and Extinction

One week prior to behavioral testing, Miniscope mice were habituated to the Miniscope for 2 days in 10 min sessions in the home cage. All mice underwent behavioral training and testing in Med Associates fear conditioning boxes for five days. In Context A, Med Associates chambers were equipped with smooth white floor inserts and cleaned with ethanol to provide a unique olfactory, tactile, and visual context. In Context B, the shock grid floor was exposed, mouse bedding was placed in a tray under the floor, and chambers were cleaned with Clidox. The five days of behavioral testing consisted of *Habituation* (Hab), *Acquisition* (Acq), and *Extinction 1-3* (Ext1-3). Hab and Ext1-3 took place in Context A, and *Acquisition* took place in Context B. The CS consisted of a 4kHz, 75dB tone delivered in 25, 200ms pips at 1Hz. During *Acquisition*, the CS was followed by a 20sec trace period preceding a 1mA, 2sec shock. On all other days, the shock was omitted. *Habituation* and *Acquisition* consisted of 8 trials, with jittered ITIs of 60±10sec. During *Extinction 1-3*, there were 6 trials per session. 30 minutes prior to *Extinction 1*, mice were injected with 1mg/kg psilocybin, contributed by the Elizabeth Heller Laboratory at the University of Pennsylvania, or saline. Mice were excluded from the study if they froze ≤20% of the time during *Acquisition* or ≤10% of the time during the first half of Extinction 1 (n=23 mice, Supplementary Fig. 1D). Two mice were excluded due to excessive barbering in the home-cage during the days of the experiment.

For Miniscope studies, a 2”-diameter hole was drilled in the top of a Med Associates box to feed the cables through. During the sessions, recordings were remotely controlled and streamed to a laptop for live monitoring. Recordings were made at LED power (0.7-1.5mW), gain (1.0-3.0), and focus (0-300μm) settings deemed appropriate for each mouse and kept as consistent between recording days as possible.

For the non-shock control condition, Miniscope-implanted mice underwent an identical protocol, except for the total omission of the shock.

#### Surgery

For Miniscope studies, all mice were unilaterally injected with 800nL of AAV9-syn-GCaMP8m-WPRE at a titer of 1.2e12 (Addgene virus #162375) in the RSC. RSC coordinates were chosen from past studies: -2.25 AP, +0.3ML,-0.8 DV. Mice were anesthetized with isoflurane. Hair was removed with Nair, and the skin sterilized with Betadine and ethanol. An incision was made with scissors along the scalp. Tissue was cleared from the skull surface using an air blast. The skull was leveled such that the Bregma-Lambda and ML DV difference was within ±0.1mm. A craniotomy was made at the chosen coordinates with a dental drill. A needle was lowered to the target coordinates through the craniotomy and virus infused at 100nL/min. The needle was left in the brain 10mins after infusion before being slowly withdrawn. The incision was sutured, and the animal was administered Meloxicam before being placed under a heat lamp for recovery.

Miniscope implantation surgeries subsequently followed the same protocol until the craniotomy step. A 1mm craniotomy was made by slowly widening the craniotomy with the dental drill. Dura was peeled back using microscissors, sharp forceps, and curved forceps. The craniotomy was regularly flushed with saline, and gel foam was applied to absorb blood. An Inscopix Pro-View Integrated GRIN lens and baseplate system was attached to the Miniscope and a stereotax. Using the Inscopix stereotax attachment, the lens was slowly lowered into a position over the injection site. The final DV coordinate was determined by assessing the view through the Miniscope stream. If tissue architecture could be observed in full focus with light fluctuations associated with RSC slow oscillatory activity under anesthesia, the lens was implanted at that coordinate (-0.6 to -0.3DV). The GRIN lens + baseplate system was secured to the skull with Metabond and then dental cement. After surgery, mice were singly housed and injected with Meloxicam for three consecutive days during recovery.

#### Miniscope validation

Before admission to the experiment, the miniscope was magnetically attached to each animal’s implant for habituation and streamed using the Inscopix Data Acquisition Software. If many cells could be observed during spontaneous behavior in the home cage, the mouse was admitted. If only a few cells were visible, the session was recorded and analyzed in the Inscopix Data Processing Software (IDPS) to determine the number of observable cells. If an animal had >20 identifiable cells, they were admitted into the study. Others were euthanized.

#### Histology

Animals were perfused with 10% formalin and brains dissected. Brains were stored in formalin solution for 24 hours before being transferred to 30% sucrose. Brains were sectioned at 50μm on a cryostat and stored in PBS. RSC sections were stained with DAPI and sections from -2.18AP to -2.88AP were mounted on slides. The section with the deepest and widest GRIN lens track was designated as the coordinate of implant.

### Analysis Methods

#### Behavioral

Behavior was recorded by Basler cameras into Pylon Viewer at 15Hz. Videos were then processed in the open source ezTrack Jupyter Notebook. The algorithm was calibrated to the standard light fluctuations in the empty chambers and the empty chambers with the Miniscope wire dangling in them for each respective study. A freezing threshold was determined in terms of number of pixels changed/frame by visually validating portions of videos classified as “Freezing” or “Moving” by the algorithm. In general, a freezing threshold of 50-200pixels/frame was used in non-Miniscope studies, whereas a threshold of 300pixels/frame was used in all Miniscope animals, necessitated by movements of the Miniscope wire. An animal was only classified as “Freezing” if the pixels/frame remained below threshold for at least 1sec, or 15 frames. Freezing status per frame was exported in a CSV file and post-processed in Matlab to calculate % freezing windows of time. Freezing plotted here is % freezing during the trace period, as this is the interval of time invoking the RSC for fear and extinction encoding and retrieval. Freezing videos were aligned to trial times by beginning analysis at the first frame of the red light in the Med Associates boxes switching on, indicating session start. Although tone delivery times were pseudo-random with respect to the animals, they were hard-coded by the experimenter, so analysis alignment to session start was sufficient to align video to tone.

#### Calcium imaging pre-processing

Videos were downloaded from the Inscopix Data Acquisition Box and uploaded to the Inscopix Data Processing Software (IDPS). Videos were spatially downsampled by a factor of 4 and spatial bandpass filtered between 0.005 and 0.500. Videos were then motion corrected with respect to their mean frame. Cells were identified and extracted using CNFM-E (default parameters in the Inscopix implementation of CNMF-E, except the minimum in-line pixel correlation = 0.7 and minimum signal to noise ratio = 7.0) and second-order deconvolved using SCS. Videos across 5 days of behavioral training were longitudinally registered in IDPS (minimum normalized cross-correlation = 0.1). Only cells registered on all 5 days were considered for further analysis.

#### Calcium imaging post-processing

Most subsequent analysis was performed in custom Matlab scripts, available in the associated GitHub. Deconvolved calcium traces of cells from each session were aligned according to their global cell index determined in longitudinal registration. As the window considered for each trial included a 10sec baseline period, a 25sec stim period, a 20 sec trace period, a 2 sec shock/omission period, and 3 sec after, each trial was 60sec. Neural activity was therefore summed within 1 sec time windows. Miniscope recordings were started exactly 30.00sec before behavioral session start, and this information was used to align data to behavior and neural data. To determine whether a cell was stimuli-, trace-, and/or shock-responsive, their baseline period activity was compared to their activity during the period of interest by permutation test in 1000 iterations. The proportion of stim/trace/shock-responsive cells compared to all longitudinally registered cells recorded within an animal was calculated for each session and compared between groups and over time with a Two-Way RM ANOVA. When all recorded cells were considered, proportions of these cells compared to the total population were compared within session across groups with a Two-Way ANOVA. A cell was considered stable if it was responsive in both the first and last two trials of a given session; recruited if it was not responsive in the first two but responsive in the last two; and suppressed if the opposite was true. Pearson’s rho was used to calculate the correlations between the proportion of these cells and total % freezing in a session. To calculate the change of activity in groups of neurons between sessions, activity was zscored to traces recorded in *Acquisition* and compared between groups and over time with a Two-Way RM ANOVA. To calculate overlaps between ensembles of neurons, ensembles were identified by TCA (described below) in each animal. Whether these overlaps were small or large was determined by a Wilcoxon ranksum test comparing the median of each overlap in each group with a 50% threshold. If an ensemble shared significantly <50% of neurons with another, this was considered a small overlap. All statistics were calculated in Prism.

#### Freezing encoding

To calculate freezing encoding in single longitudinally registered neurons, neural traces were downsampled from 20 to 15Hz and aligned to a 15Hz binary freezing trace. A binomial GLM was trained on half of the data from each session and evaluated on the other half to generate auROCs. The mean auROC of all neurons in a mouse in each session was reported in the main text. To determine the population encoding of freezing in the RSC, 15Hz activity of each recorded cell was normalized to its maximum. 15Hz PSTHs from 2 seconds before to 2 seconds freezing onset or freezing offset of all cells from each day were averaged over trials and projected into the same principal component space, unique to each animal. The Euclidean distance was calculated between these trajectories at each timepoint and averaged across groups for Fig. 3D and then averaged over time for Fig. 3E. To determine whether changes in freezing encoding in PC space were observable at the single cell level, d-prime was calculated for each neuron over the same motion and freezing time windows used to produce the trajectories. Where *a* = activity:

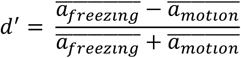

Single cell freezing discrimination for each animal was reported as the median of the absolute values of d-prime. These outcomes were compared within sessions between treatment and extinction rate groups with a Two-Way ANOVA.

#### TCA

To perform TCA, post-processed calcium imaging data was arranged into tensors *t*_sec_ x *c*_cells_ x *T*_trials_ in size for each animal, exported as a Matlab structure, and imported into Spyder where we employed the TensorTools package developed by Williams et al., 2018. To determine the appropriate model rank empirically, TCA was first run on the pooled and aligned tensors from all mice in each treatment group, and model reconstruction error and similarity were plotted as a function of increasing model rank. The elbow method revealed that models of rank 5 were most appropriate for subsequent analysis (Reconstruction error = 0.615; Model similarity between four iterations = 1). Models of rank 5 were then generated for each animal.

To measure the dominance of each of the 5 components during a given trial, the relative strength of a given component was measured as the fraction of the total trial weights at that time assigned to that given component. This measure functions to assess how dominant this component is over others at a certain time. Linear regression was used to determine the relationship between component-dominance and behavior over time.

To determine total extent of session discriminability of TCA-identified components, pooled TCA models were generated 100 times for each group and compared to models generated on 100 shuffled datasets from the same groups using unpaired t-tests. Neurons were randomly shuffled at each timepoint to preserve the within- and across-trial temporal structure of the data, controlling for changes in recording quality across days. To compare across groups, neurons were randomly subsampled in each iteration of TCA to control for effects of the number of cells on session discriminability.

As TCA also assigns weights to each neuron in each component, we found we could use this information to identify ensembles of neurons driving each component.

#### Identifying neural ensembles

TCA returns neuron factor loadings signifying the relative weight of each neuron in each component. However, the absolute values of these weights are influenced by the size of the data tensor across all three dimensions. To determine the neuron factor loading or weight above which a neuron would be contributing to a component greater than by chance, simulated data tensors were generated for each animal populated with identically behaving neurons. For animal *a* with *c* longitudinally recorded neurons, given a constant experimental structure of T = 34 total trials with t = 60 sec per trial, a tensor of 60 sec x c_a_ x 34 trials was generated and TCA iterated 100x. We chose a threshold of w=1.0 as the median and mean of the null distribution of the factor loading threshold were greater than 1.0 and less than 1.1. Primary outcomes (See **Fig. 5B-D, Supplementary Fig. 6**) were re-calculated using various factor thresholds to verify that results with a threshold w=1.0 are robust to threshold choice. Thus, *Acq-*dominant ensemble, for instance, was therefore comprised of neurons with w>1 in the *Acq-*dominant component determined by the strength metric described above.

#### Multiple regression

To determine the predictive power of the observed changes in neural ensemble activity over freezing in late Extinction 3, mean % trace period freezing, mean change in activity from Acquisition of the *Acq-*dominant ensemble during Extinction 1 and of the *Ext3-*dominant ensemble during Extinction 3 were all respectively z-scored within treatment condition to normalize the distributions and regressed. Adjusted R^2^, p values, coefficients, and their 95% confidence intervals are reported.

#### Fisher linear discriminant analysis

A Fisher decoder was trained in Matlab on one of seven predictors: the mean activity of the *Acq-Only, Ext1-Only, Ext3-Only, Acq/Ext1, Acq/Ext3, Ext1/Ext3*, or *Acq/Ext1/Ext3* ensembles over all timepoints in each session. Class labels were “Responders,” “Non-responders,” or “Rapid” mice. Fisher decoders were trained to distinguish between data from two of the class labels to determine how similar or different the ensembles between pairs of groups behaved. Fisher decoders were trained on a randomly selected 50% of the data and evaluated on the other 50% over 100 iterations. As a control, class labels were randomly shuffled, and model performance was evaluated on the shuffled data. If the accuracies of the decoders generated by a given ensemble’s activity overlapped with the distribution of accuracies when evaluated on shuffled data, it was classified as failing to predict responder status or treatment. If not, then this ensemble was classified as predictive with respect to the given distinction. To validate the ability to distinguish all three classes based on these ensembles was verified with three-way Fisher decoders in Supplementary Fig. 4.

#### Computational model

A linear non-linear firing rate model was composed of a hypothesized RSC microcircuit comprised of *Acq-* and *Ext3*-dominant neurons obeying the following system of equations based on the cortical circuit model proposed by Park and Geffen, 2022^80^. To evaluate the activity at time t in neuron I, take the following quantities as time t-1:

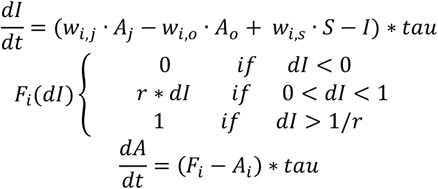

Where *I* is the synaptic input to neuron *i, w*_*i,j*_ is the weight of input from neuron *j* in the same ensemble, *w*_*i,o*_ is the weight of input from neuron *o* in the opposing ensemble, *A* is the presynaptic neuron’s activity, *w*_*i,s*_ is the neuron’s selectivity matrix for the stimulus, *S* is the stimulus matrix for tone, trace, and shock/omission inputs, and tau is the time constant (1ms where *dt* is 1s). *F* is the function describing the nonlinear part of neural activation, such that subthreshold inputs are scaled by a factor *r* = 3. The transformed synaptic input F is then added to the neuron’s activity *A*.

Each ensemble was modeled with 10 recurrently connected units with either selective or mixed-selective response properties mimicking the real data that mutually inhibit neurons of the opposite ensemble. (*Acq-*dominant ensemble: 80% shock-responsive, 20% tone responsive, 20% trace responsive; *Ext3-*dominant ensemble: 50% shock-responsive, 30% tone responsive, 40% trace responsive). Intra-ensemble weights underwent excitatory plasticity according to a Hebbian learning rule of where the learning rate alpha = 0.1. Psilocybin was simulated only as 100 increasing amounts of direct synaptic inhibition ranging between 0 and 1.

The model was trained on an identical TFC acquisition and extinction task as the mice, with tone and trace delivery simulated as of 0.1 to their responsive units. Shock or shock omission were modeled as inputs to shock or omission responsive neurons as an input of 1 or -1 respectively, and omission-responsive Ext3-dominant neurons weighted this input with *w*=-1 to flip the sign. The difference in magnitude between shock and tone or trace inputs is intended to reflect their differing salience. The magnitude of the shock-omission input decreased linearly with each trial after Extinction 1, intended to represent reduced salience of the absence of shock over time.

To test the model, its structure was systematically varied to challenge the following underlying assumptions: 1) to test whether the neural populations are excitatory, the weight matrices between neurons of the same ensemble were set to zero, one by one and then together; 2) to test whether the neural populations were mutually inhibitory, the input weights from one ensemble to the other were set to zero, one by one and then together; 3) to test whether shock-omission sensitivity was required, the weight of shock omission inputs to the *Ext3-*dominant ensemble was set to zero.

To test whether inhibition of the *Acq*-dominant ensemble was sufficient to explain our empirical results, psilocybin was simulated as *P* = 101 evenly incremented values of 0 to 100 and subtracted directly from the synaptic input *dI* as *P*tau* for the whole Extinction 1 epoch of training.

To determine the fit between the output activities of the model and the real data, the slopes of their linear parts were calculated using the line equation:

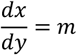

MSE between the real data and each slope was then calculated. To calculate simulated freezing in the model, activities were plugged into the equation yielded from the multiple regression of z-score % freezing on z-score activity of *Acq-*dominant neurons during Extinction 1 and *Ext3-*dominant neurons during Extinction 3:

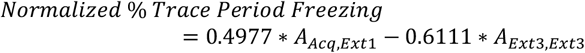

## Acknowledgements

This work was funded by NIH NIGMS DP2GM140923 awarded to G.C. We thank the University Laboratory Animal Resources (ULAR) group at the University of Pennsylvania for assistance with rodent husbandry and veterinary support, including all faculty stationed at both the Translational Research Laboratory. Thanks to Dr. Maria Geffen (Penn) for advice on model construction and Stephen Wisser (Penn) for assistance running behavioral experiments. We would also like to thank other members of the Corder Lab, Adrienne Jo (Penn) and Raquel Aiada Sandoval Ortega (Penn), and for critical discussions and advice on behavioral analysis, data visualization, and analysis validation. We would also like to thank Colin Mackey for assistance in customizing various python packages. Finally, we would like to thank the faculty of the Cold Spring Harbor Laboratory course in Neural Data Analysis for critical input on analysis approach.

## Author contributions

S.R. and G.C. conceptualized and planned out the study. E.H. provided key resources including psilocybin and assisted with experimental design and behavioral analysis. S.R. performed all data collection, analysis, and writing. G.C. acquired funding, performed data visualization along with S.R., and edited and revised manuscript.

## Supplementary Materials

**Supplementary Figure 1.**
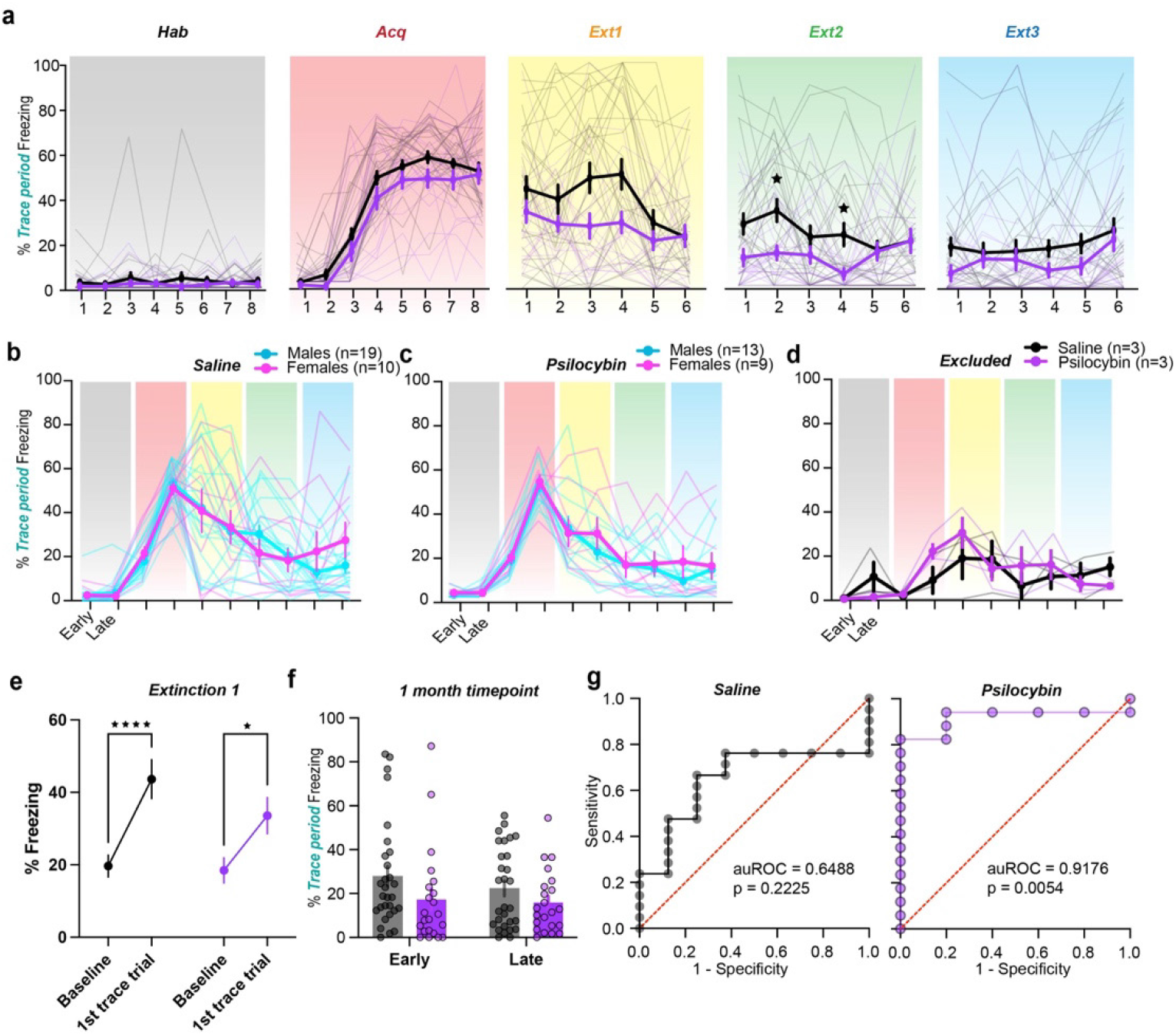
│ Effects of psilocybin on trace fear extinction in males and females. **a**. Trial by trial freezing of saline- and psilocybin-administered mice. Two-Way RM ANOVA with Sidak multiple comparisons correction. (Supp. Table 1, rows 102-106) **b**. Half-session freezing by sex of saline-administered animals. Two-Way RM ANOVA with Sidak multiple comparisons correction. (Supp. Table 1, row 107) **c**. Same as B) in psilocybin-administered animals. Two-Way RM ANOVA with Sidak multiple comparisons correction. (Supp. Table 1, row 108) **d**. Half-session freezing by treatment of excluded animals. Two-Way RM ANOVA with Sidak multiple comparisons correction. (Supp. Table 1, row 109) **e**. Percent freezing during the baseline vs. trace period during Extinction 1. Two-Way RM ANOVA with Sidak multiple comparisons correction. (Supp. Table 1, row 110) **f**. Percent trace-period freezing in early and late periods during an Extinction session 1 month after Extinction 3. Two-Way RM ANOVA with Sidak multiple comparisons correction. (Supp. Table 1, row 111) **g**. ROC curves from logistic regression predicting RE or SE status based on % time freezing during the first half of *Extinction 1* during acute drug treatment in saline-administered mice (left) and psilocybin administered mice (right). Right: ROC curve from logistic regression. (Supp. Table 1, rows 12, 14) * p ≤ 0.05, ** p < 0.01, *** p < 0.001, **** p < 0.0001.

**Supplementary Figure 2.**
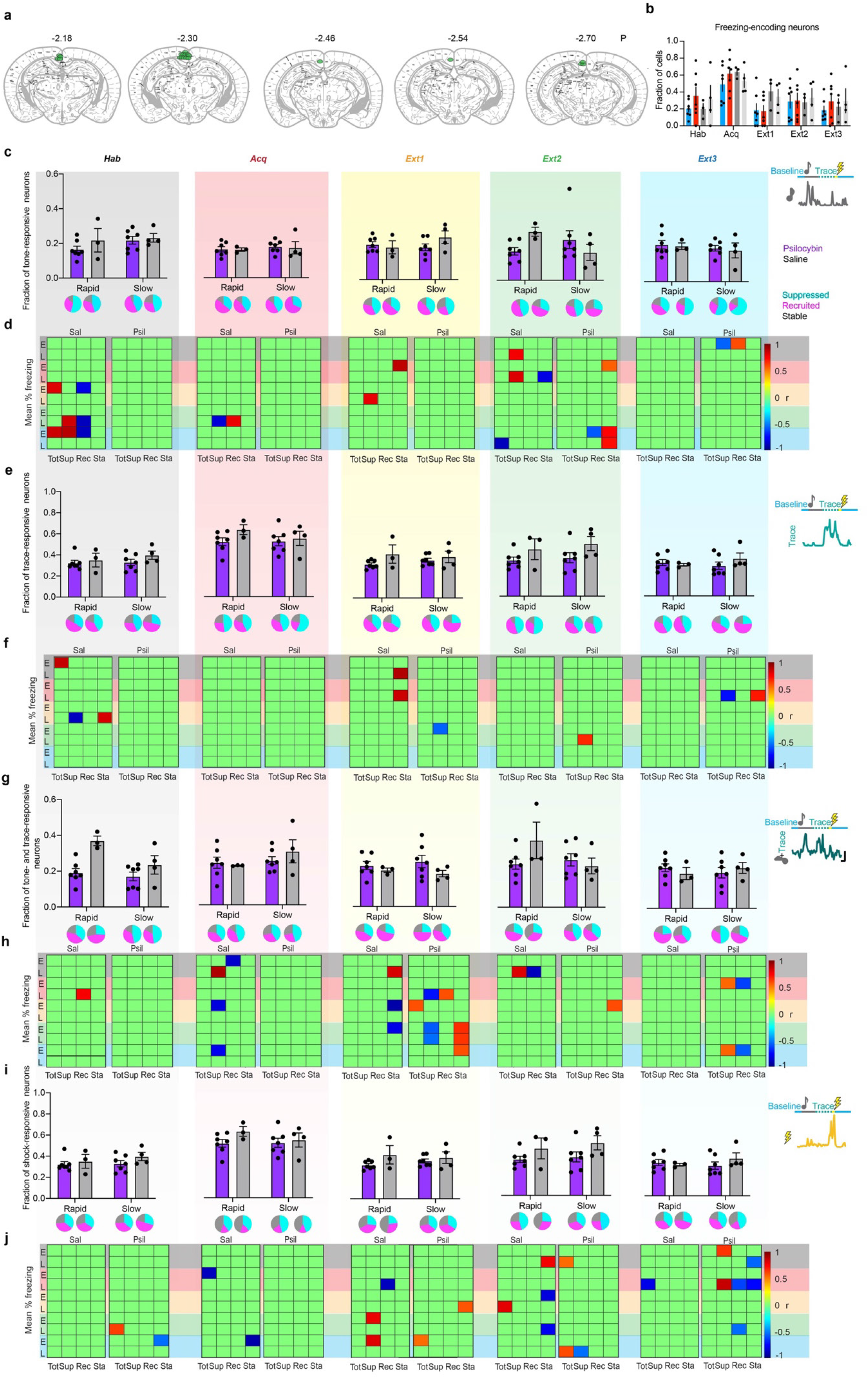
│ All RSC cells recorded. **a**. Center and bottom of implant tracts of all included mice from anterior (left) to posterior (right) granular RSC. **b**. Fraction of freezing encoding neurons on each day. Two-way RM ANOVA. (Supp. Table 1, row 112) **c**. Mean fraction of tone-responsive neurons on each day. Insets are proportions of neurons with suppressed, recruited, and stable responses. Two-Way ANOVA. (Supp. Table 1, rows 113-117) **d**. Heatmaps displaying significant correlations (Pearson’s rho) between proportions of total (Tot), suppressed (Sup), recruited (Rec), and stable (Sta) tone-responsive neurons on each day and % freezing during the early (E) and late (L) halves of each session (black rows = Hab freezing and black columns = fractions of neurons during Hab, red = Acq, yellow = Ext1, green = Ext2, blue = Ext3). **e,g,i**. Same as C for trace-, tone-and-trace, and shock-responsive neurons. Two-Way ANOVA. (Supp. Table 1, rows 118-132) **f,h,j**. Same as D for trace-, tone-and-trace, and shock-responsive neurons. Data are represented as mean ± SEM. * p ≤ 0.05, ** p < 0.01, *** p < 0.001, **** p < 0.0001.

**Supplementary Figure 3.**
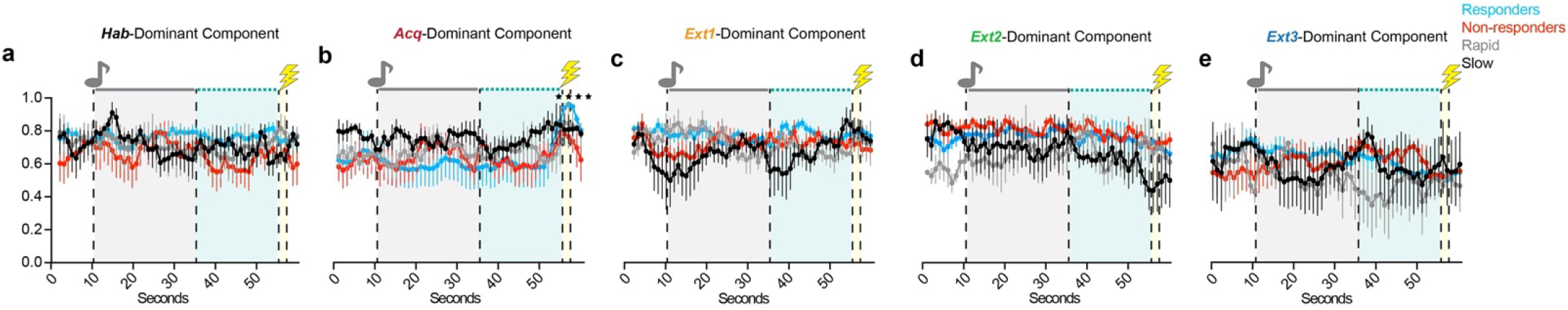
TCA factors reveal RSC dynamics modulated by session. **a**. Normalized temporal factor weights by group of the Habituation-dominant component. Two-Way RM ANOVA. (Supp. Table 1, row 133) **b**. Same as A) for the Acquisition-dominant component. Two-Way RM ANOVA. (Supp. Table 1, row 134) **c**. Same as A) for the Extinction 1-dominant component. Two-Way RM ANOVA. (Supp. Table 1, row 135) **d**. Same as A) for the Extinction 2-dominant component. Two-Way RM ANOVA. (Supp. Table 1, row 136) **e**. Same as A) for the Extinction 3-dominant component. Two-Way RM ANOVA. (Supp. Table 1, row 137) Data are represented as mean ± SEM. * p ≤ 0.05, ** p < 0.01, *** p < 0.001, **** p < 0.0001.

**Supplementary Figure 4.**
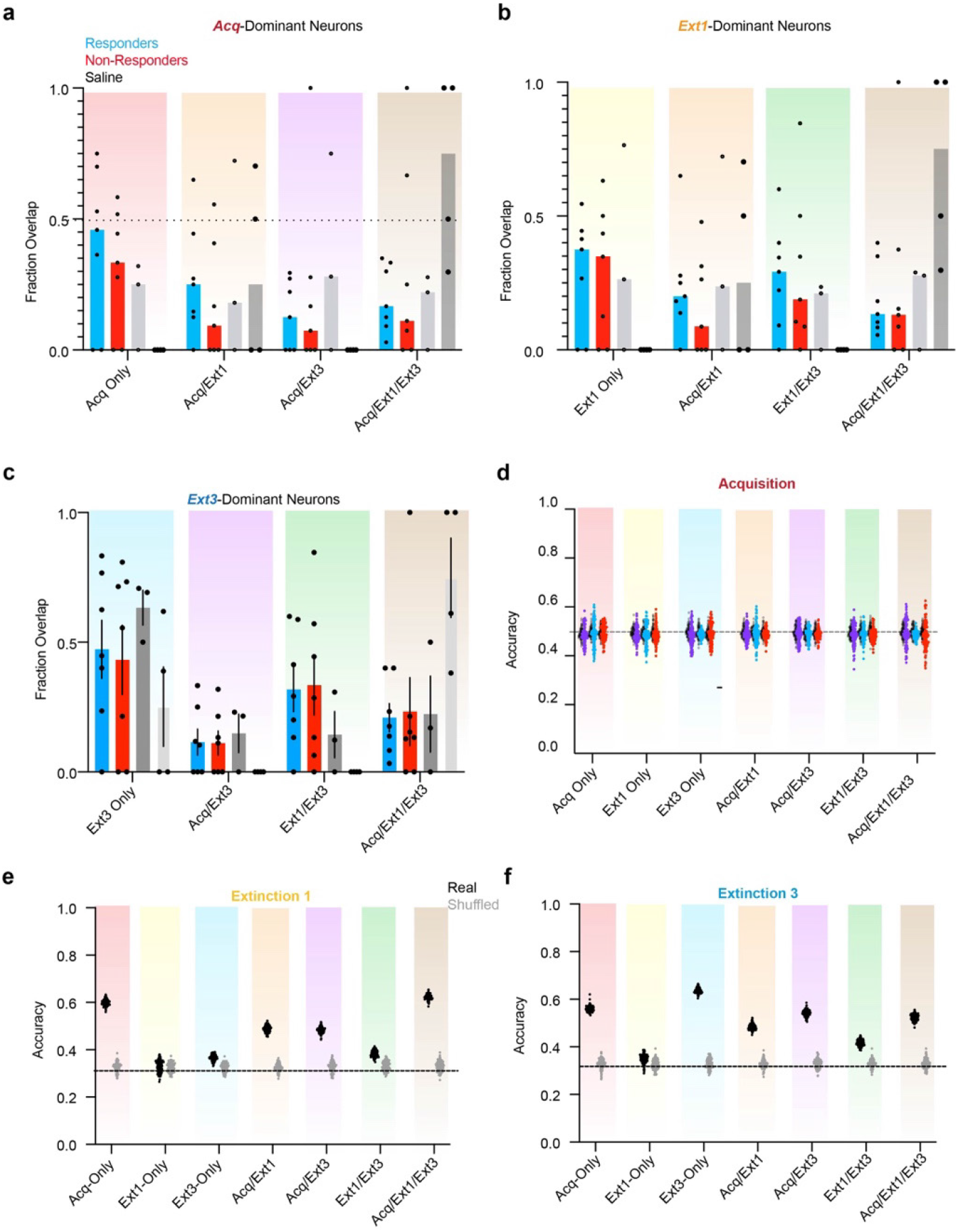
│ Psilocybin bidirectionally modulates neural ensembles driving RSC dynamics during TFC in responders. **a**. Overlaps of ensembles within individual animals comprising the mean values in Fig. 4B top. Bars are median. **b**. Same as A for Fig. 4B middle. **c**. Same as A for Fig. 4B middle. **d**. Fisher decoder performance on Acquisition activity in functionally defined ensembles of cells to distinguish responders vs. non-responders (purple), responders vs. rapid saline (blue around grey), and non-responders vs. rapid saline (red around grey). 100 iterations for each comparison. Shuffled values are behind real values. **e**. Three-way Fisher decoder performance classifying responders vs. non-responders vs. rapidly extinguishing saline mice trained on activity during Extinction 1. **f**. Same as E for Extinction 3 activity.

**Supplementary Figure 5.**
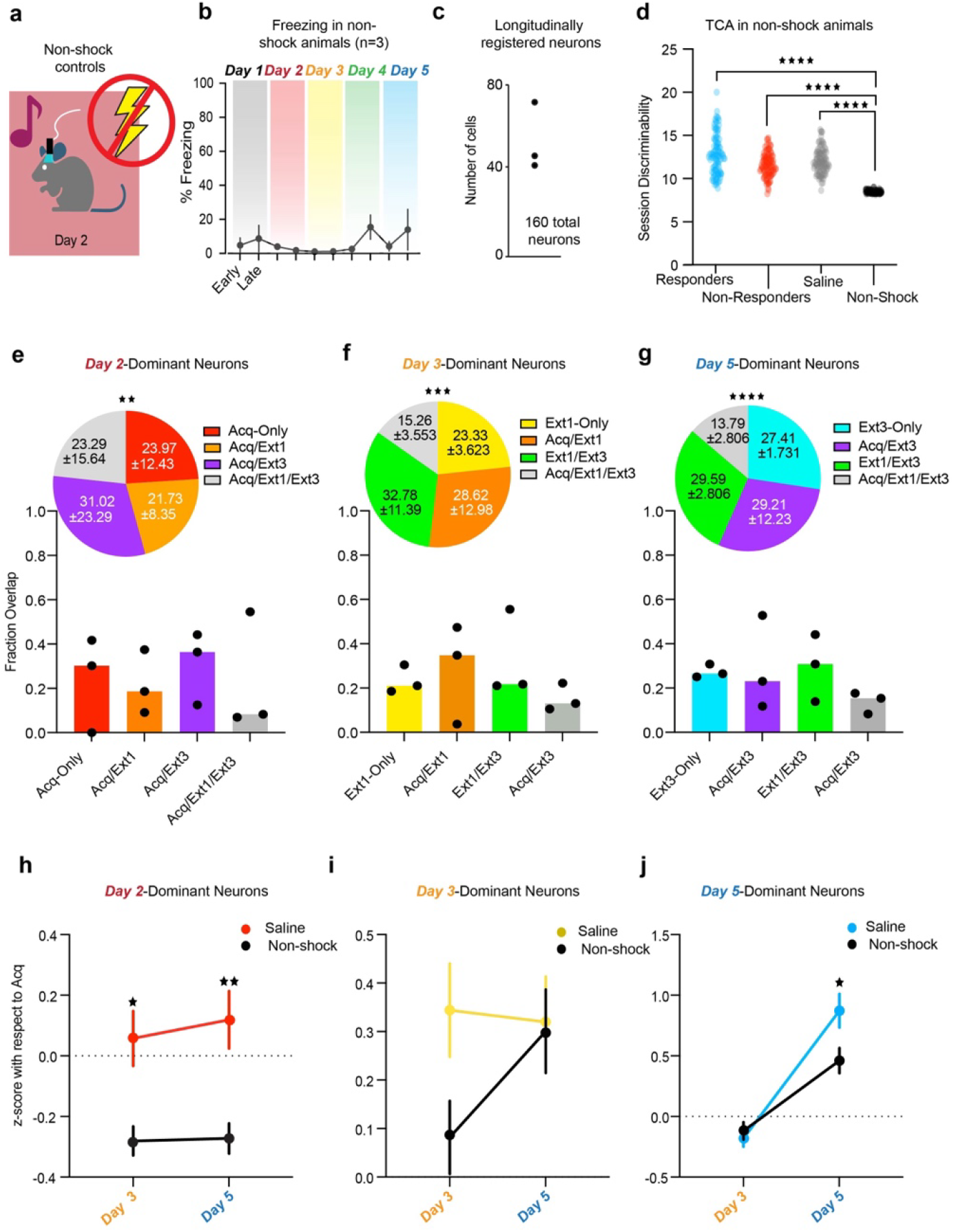
│ Non-shock controls do not exhibit conditioning-associated dynamics. **a**. Schematic of non-shock protocol. 3 Miniscope implanted mice underwent identical 5 day paradigm to all other mice, with the exception that they received no shock during Acquisition or drug treatment. **b**. Half-session freezing in non-shock mice. (Supp. Table 1, row 138). **c**. Number of longitudinally registered neurons in non-shock mice. **d**. Sum of session discriminability index. Because roughly half the number of neurons were recorded in non-shock mice as in the other two groups, pooled tensors from psilocybin responders, non-responders, and saline mice were subsampled to a different, random set of 160 neurons in each of 100 iterations of TCA. One-Way ANOVA. (Supp. Table 1, rows 139). **e**. Overlap of the *Day 2-*dominant ensemble with *Day 3*- and *Day 5*-dominant ensembles in non-shock mice. Bar graphs display the median fraction overlaps. Dots are individual animals. Insets are pie charts displaying total overlap. Stars indicate comparison to saline distribution. Chi-square. (Supp. Table 1, rows 140) **f**. Same as E for the *Day 3*-dominant ensemble. Chi-square. (Supp. Table 1, rows 141). **g**. Same as F for the *Day 5-*dominant ensemble. Chi-square. (Supp. Table 1, rows 142) **h**. Average z-score with respect to *Day 2* of *Day 2-*dominant ensemble during *Day 3* and *5* in non-shock mice (black) compared to conditioned, saline-administered mice. Two-Way RM ANOVA. (Supp. Table 1, rows 143). **I**. Same as H for the *Day 3*-dominant ensemble. (Supp. Table 1, rows 144). **j**. Same as H for the *Day 5*-dominant ensemble. (Supp. Table 1, rows 145). Data are represented as mean ± SEM. * p ≤ 0.05, ** p < 0.01, *** p < 0.001, **** p < 0.0001.

**Supplementary Figure 6.**
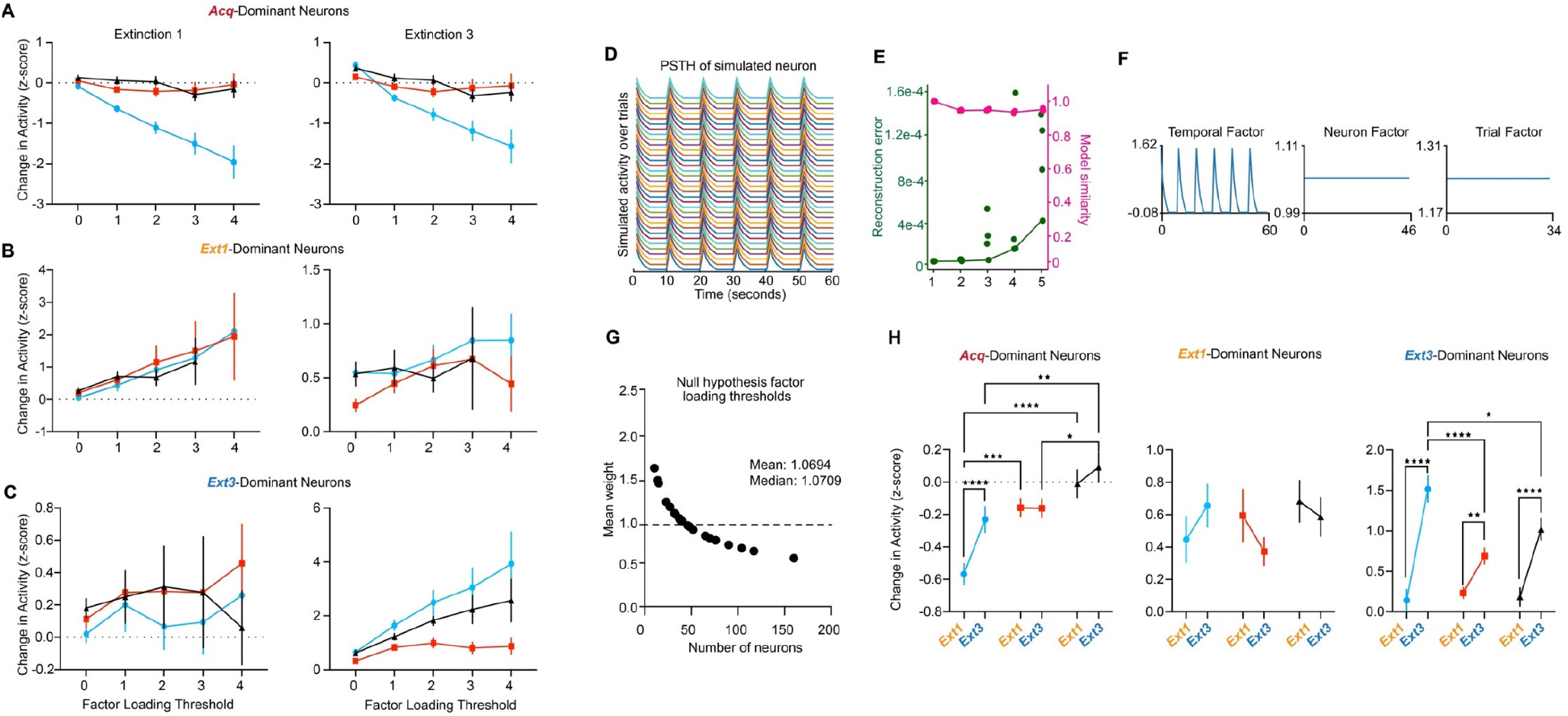
Results are robust to changes in factor loading thresholds. **a**. Change in activity in mean ± SEM from *Acquisition* in *Acq-*dominant neurons as a function of factor loading thresholds varying between *w*=0-4 during *Extinction 1* (left) and *Extinction 3* (right). **b**. Same as A) for *Ext1-*dominant neurons. **c**. Same as A) for *Ext3-* dominant neurons. **d**. PSTH of an example simulated neuron to determine the null hypothesis factor loading threshold. Tensors of t x c x T size, where c is the number of neurons recorded in a given animal, were created with identically behaving neurons to determine the factor loading threshold in a hypothetical population in which each neuron equally contributes to dynamics, or the null hypothesis factor loading threshold for that animal. **e**. Reconstruction error and model similarity of varying model ranks for populations of identical neurons. A model of rank 1 yields 0 error in this case. **f**. Representative rank 1 TCA of a simulated dataset with n=46 neurons, the median number of neurons recorded in this study. Because variances across trials and neurons were clamped at 0, only the temporal factor varies. **g**. Data in Fig. 4A plotted as a function of number of neurons recorded. Mean weight of neuron factors across 100 iterations of TCA at the number of cells recorded in each animal. **h**. Change in activity in mean ± SEM from *Acquisition* during *Extinction 1* and *3* in *Acq-*dominant (left), *Ext1*-dominant (middle), and *Ext3*-dominant (right) using ensembles determined with the null hypothesis factor loading for each animal. Two-way RM ANOVA. (Supp. Table 1, rows 88-90) . * p ≤ 0.05, ** p < 0.01, *** p < 0.001, **** p < 0.0001.

**Supplementary Figure 7.**
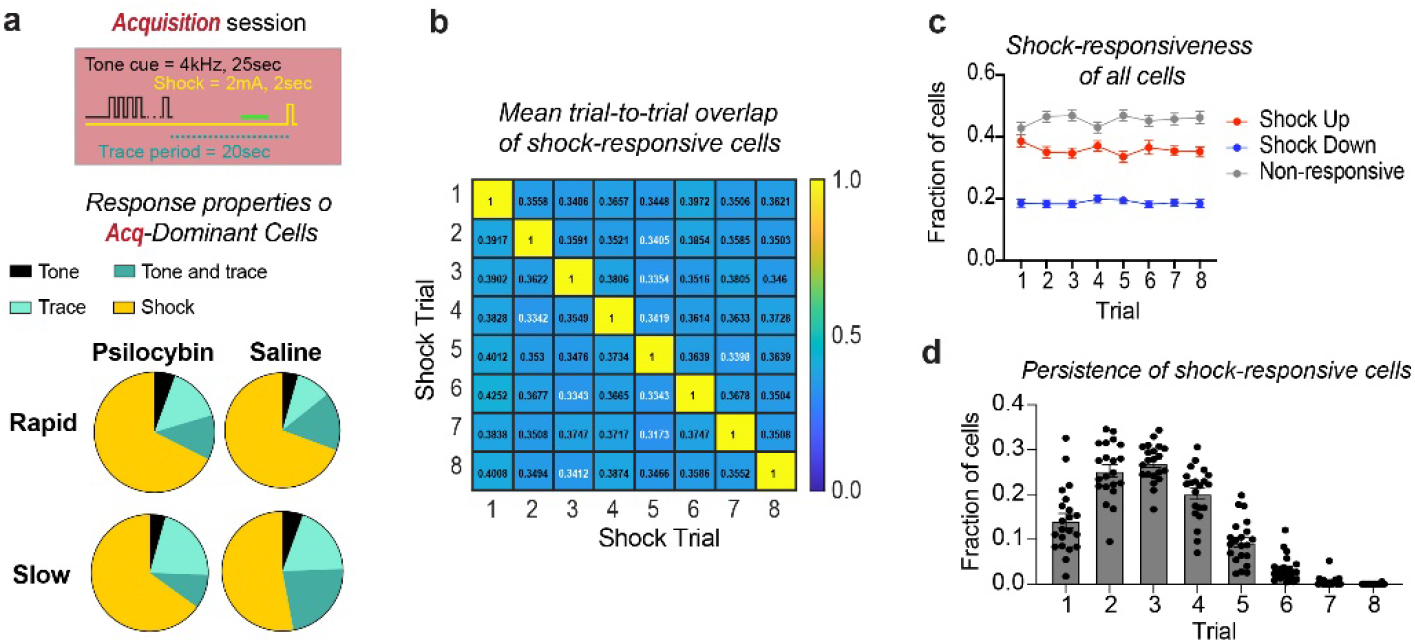
Shock-responsive neurons are unstable in the RSC. **a**. Average proportions of Acq-dominant neurons in each group that were upregulated in response to shock, trace, tone, or tone- and-trace. **b**. Heatmap of the average fraction of overlap in shock-up neurons between each trial of Acquisition. Average overlap between trials ranges from 30-45%. **c**. Fractions of shock-up, shock-down, or shock-nonresponsive neurons across all 21 mice on each trial of acquisition, determined by permutation test. **d**. Persistence of the response properties of shock-up neurons over the session. Each point y is the fraction of neurons upregulated in response to the shock for x number of trials. Data are represented as mean ± SEM over all 21 mice.

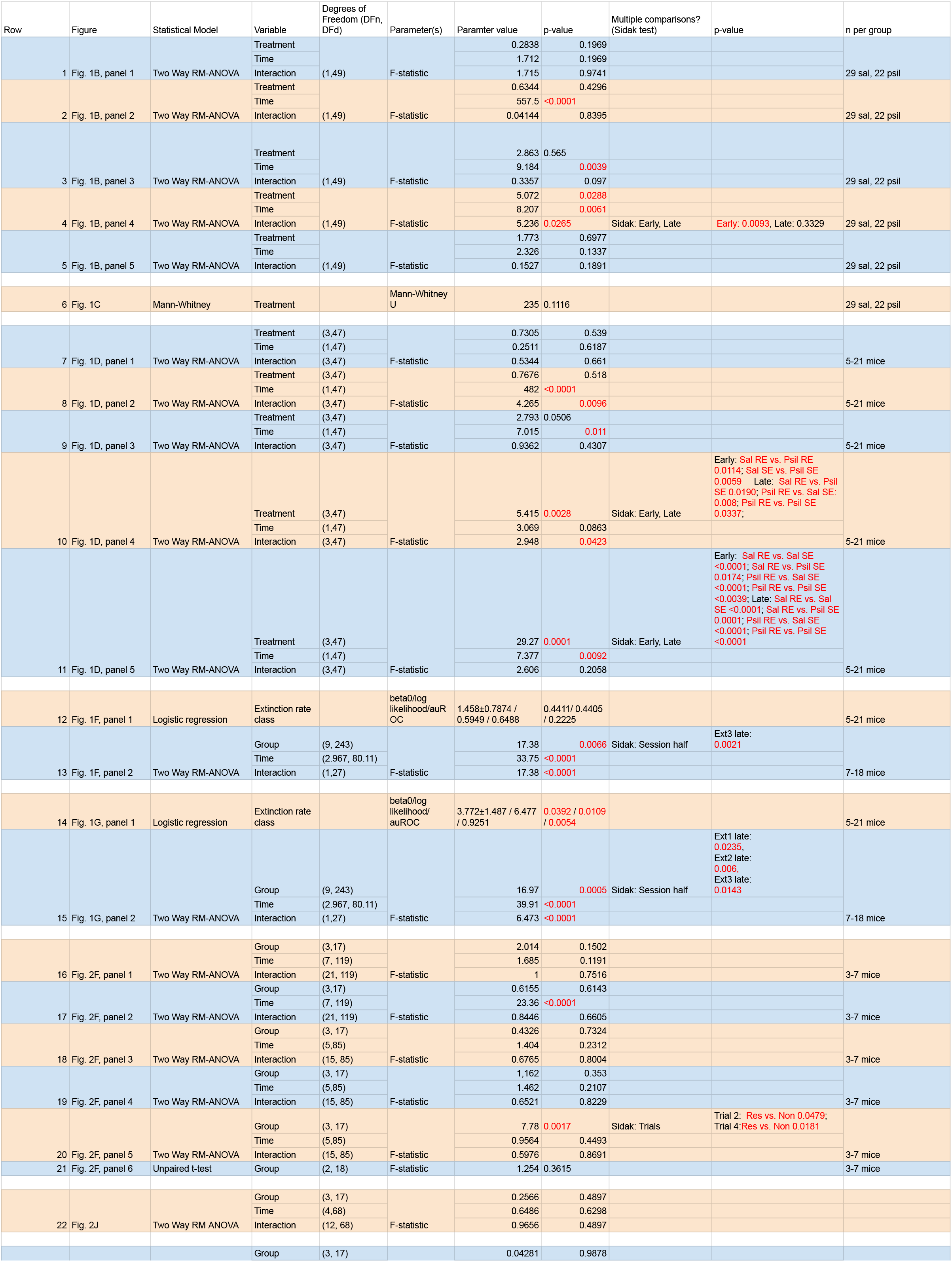
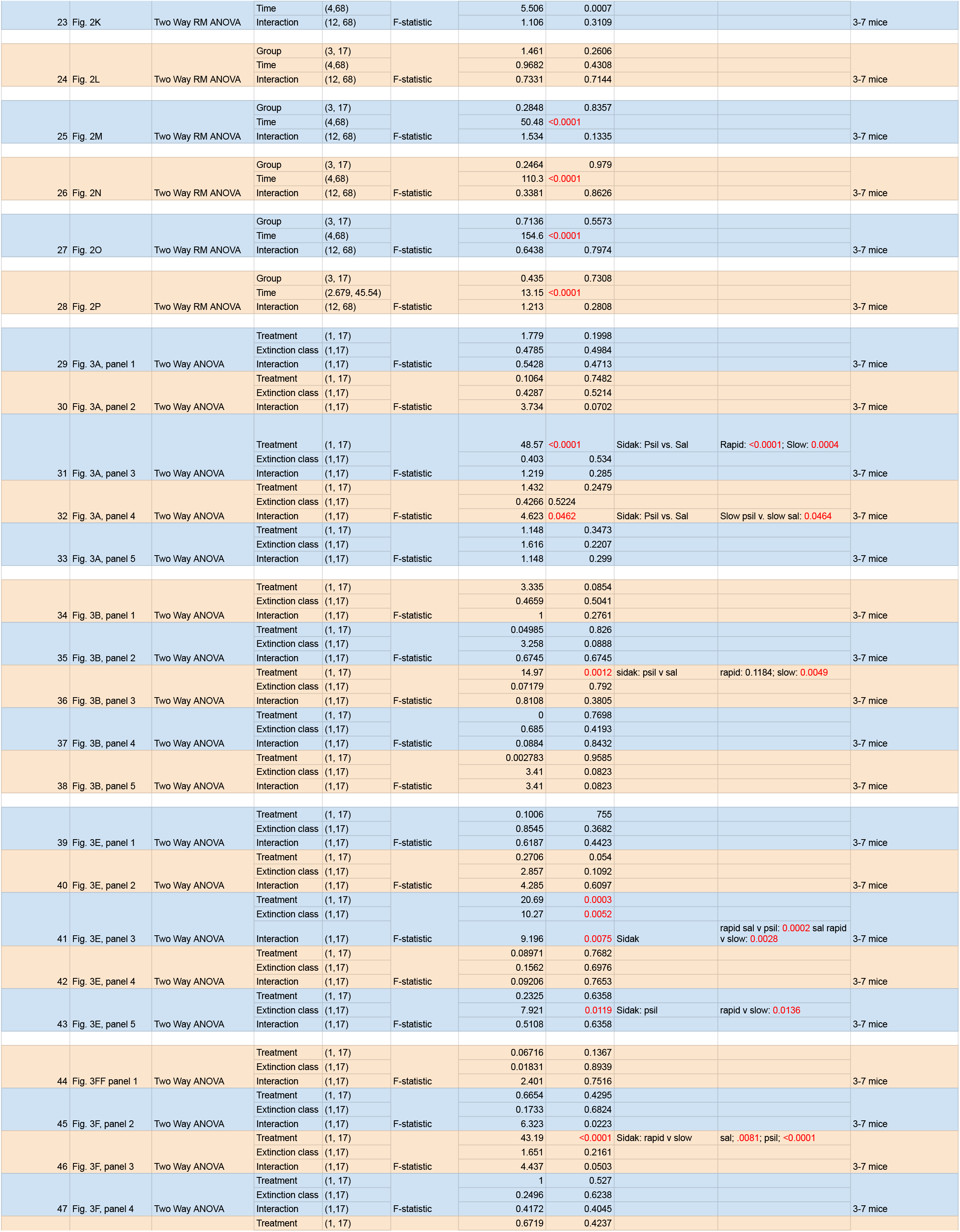
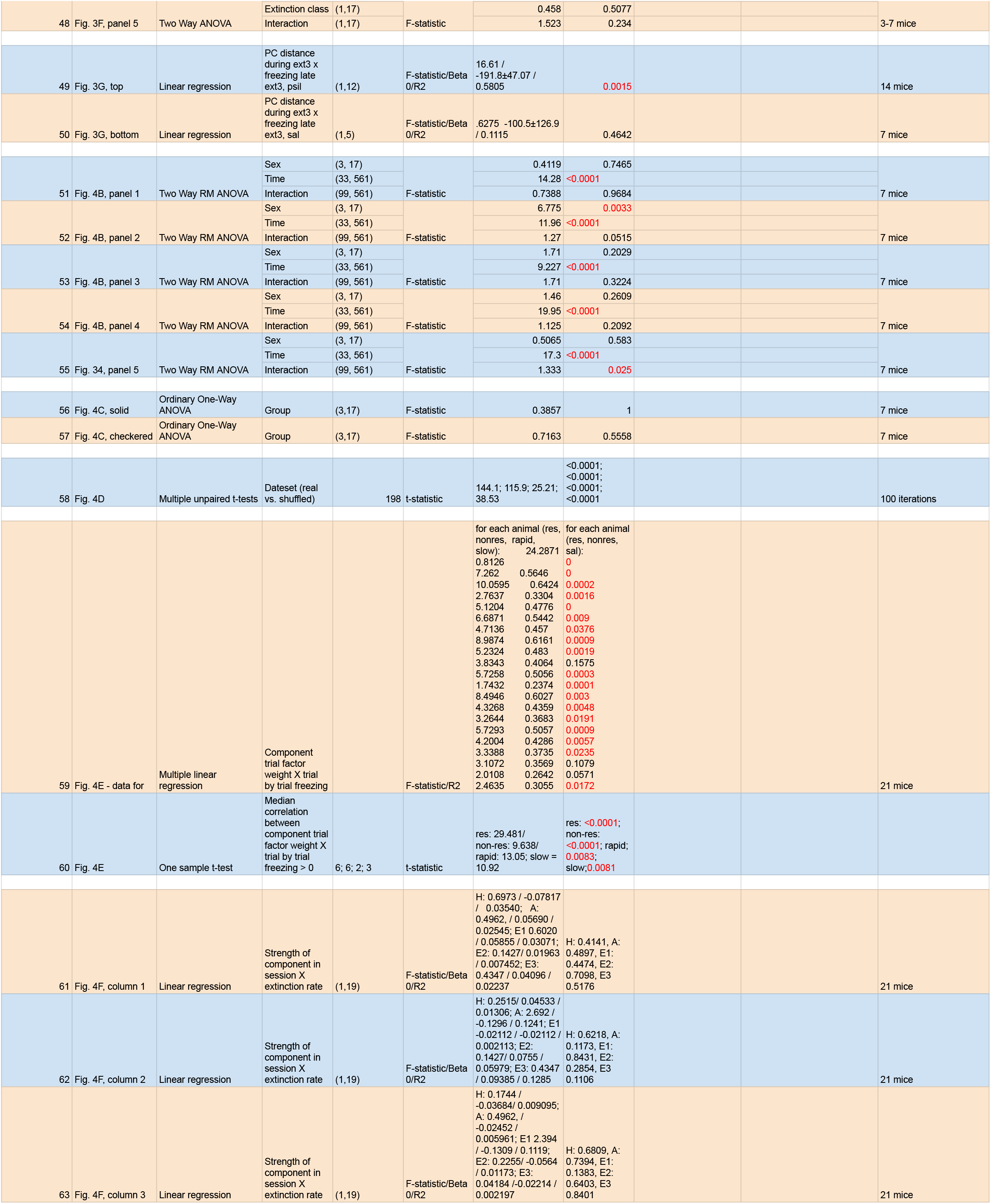
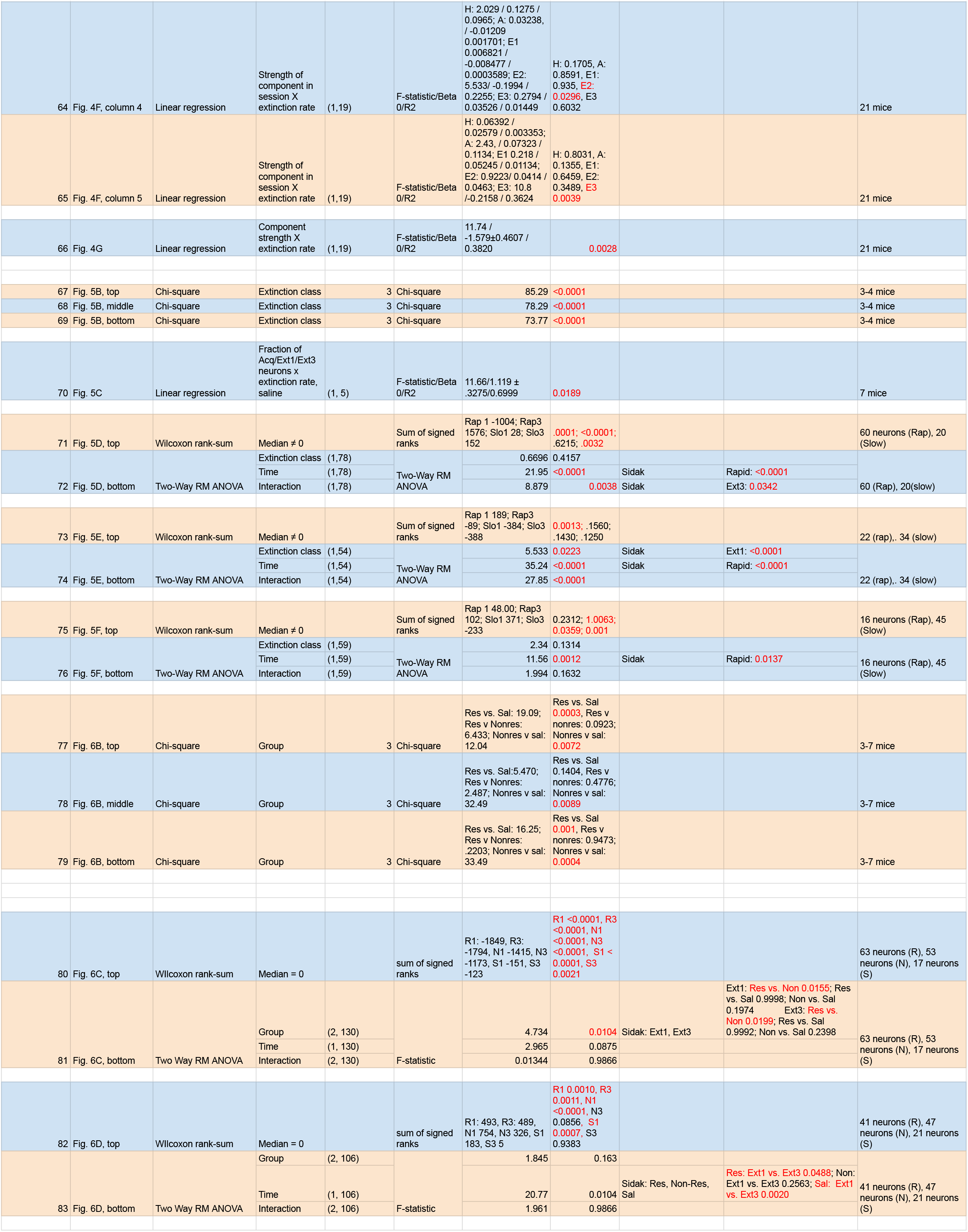
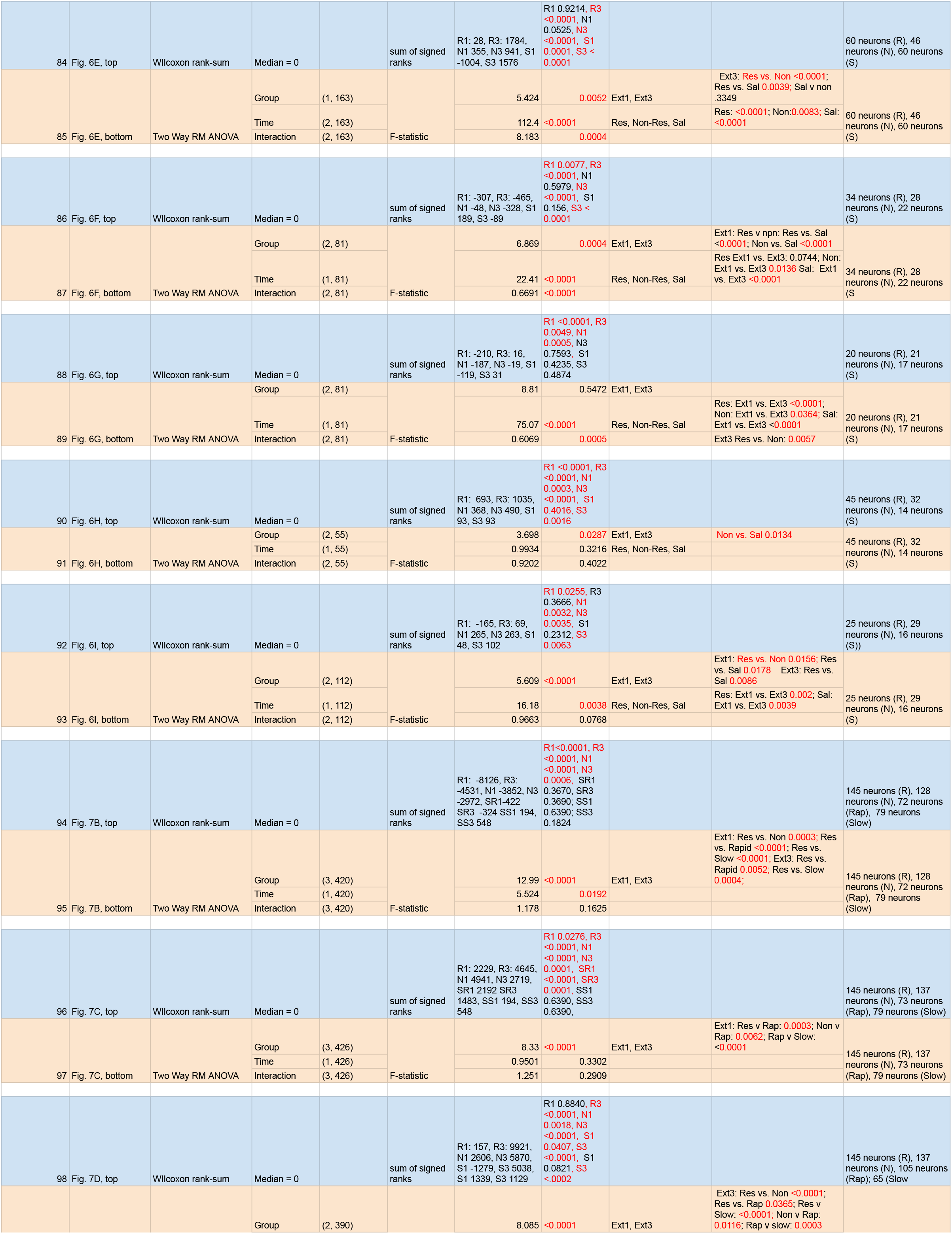
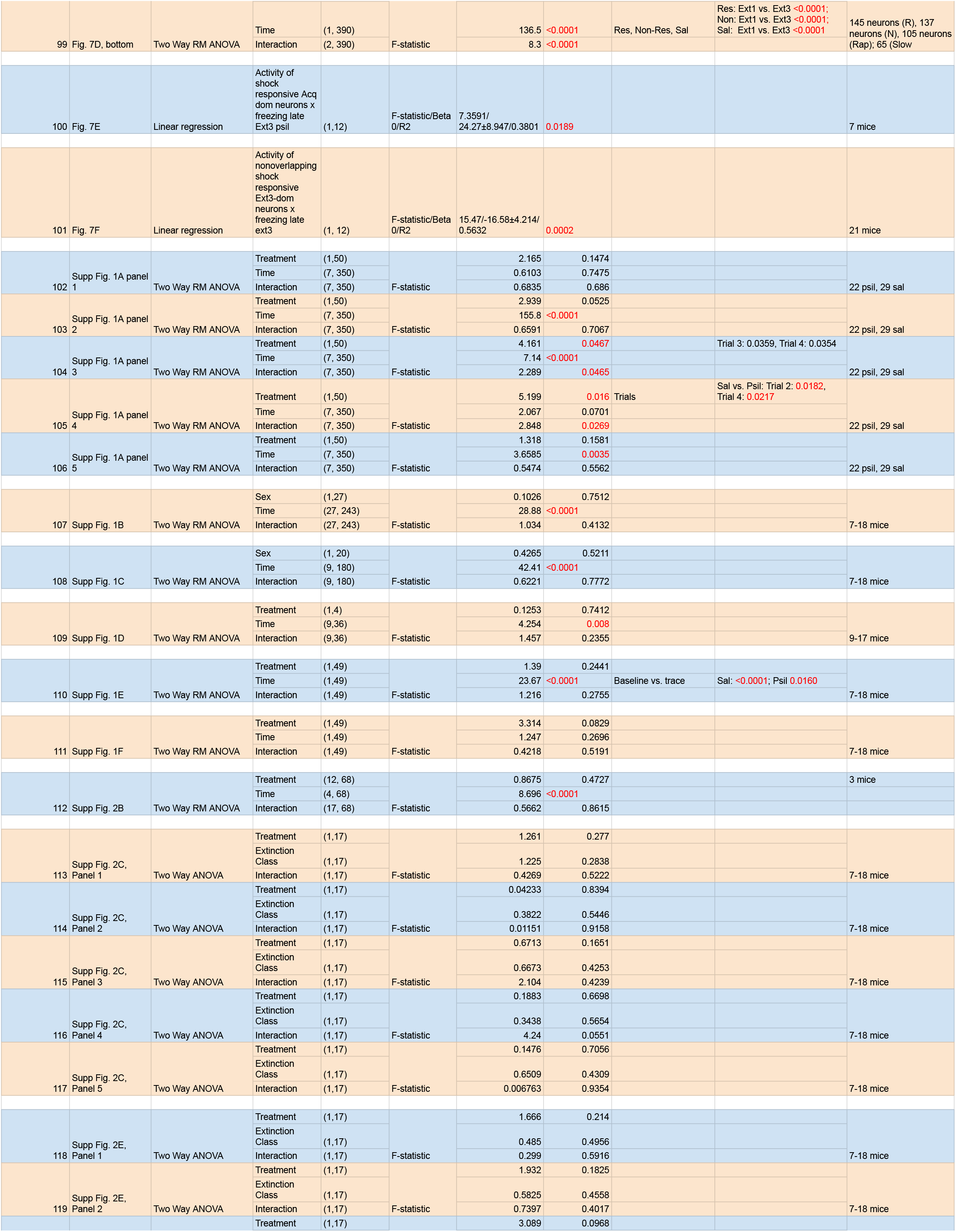
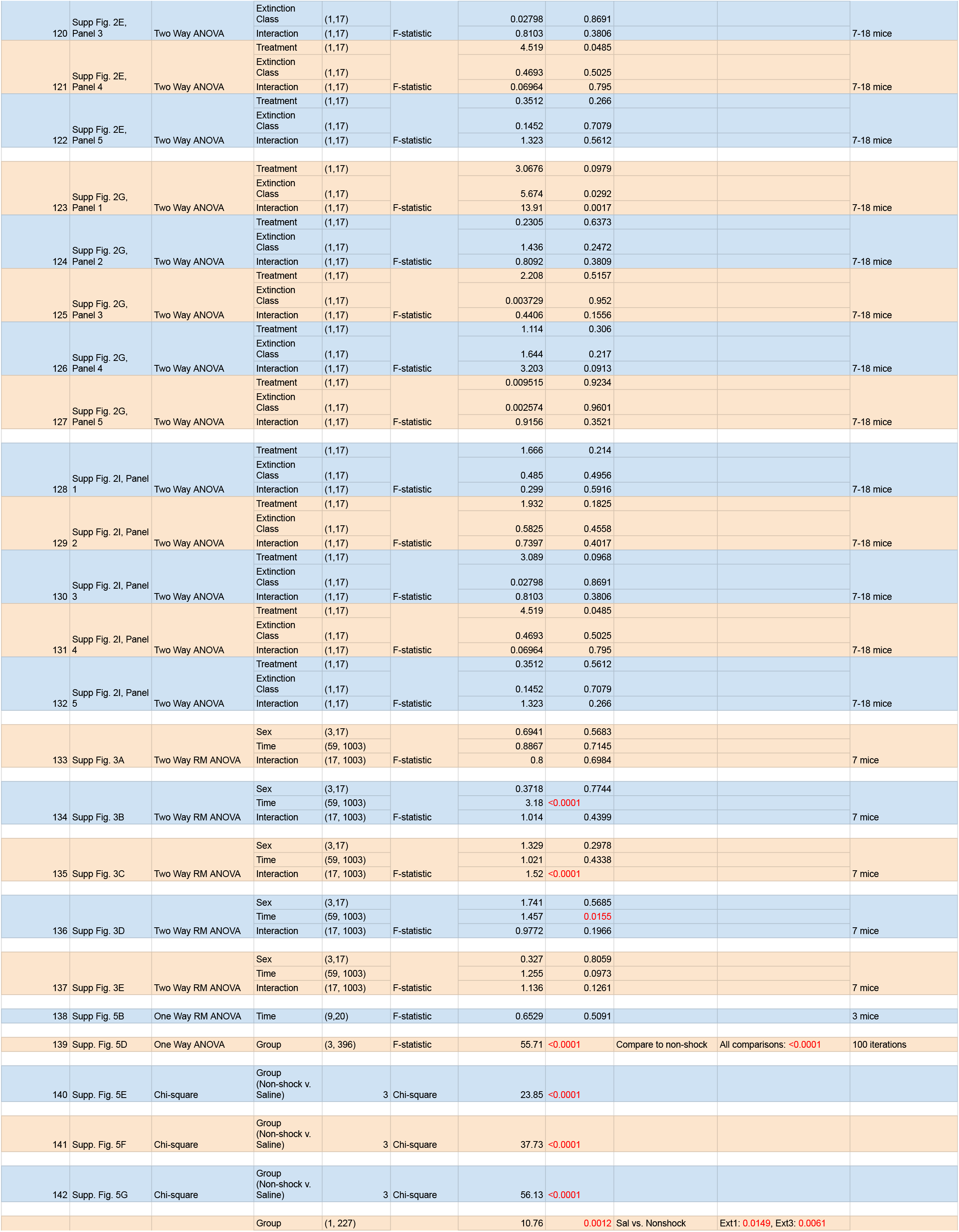
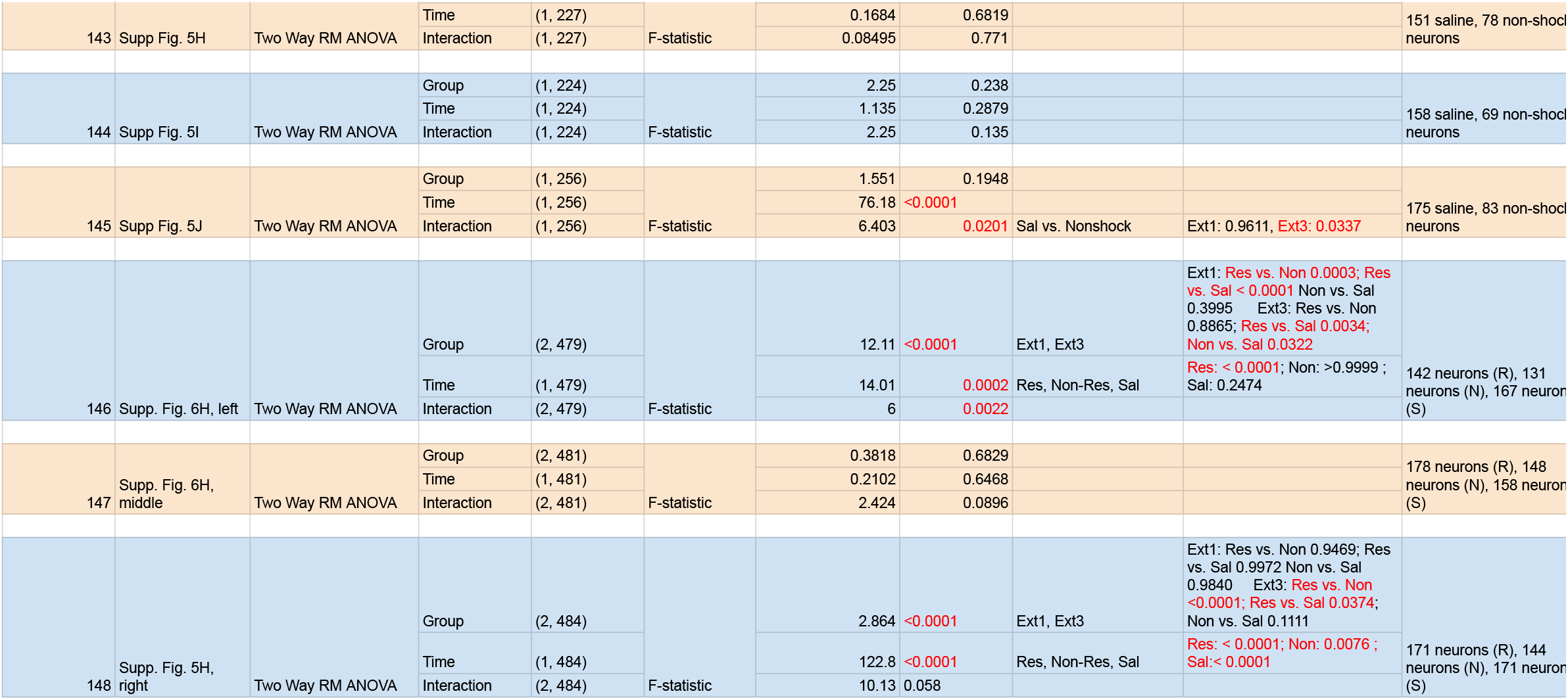

